# The Necrosis- and Ethylene-inducing peptide 1-like protein (NLP) gene family of the plant pathogen *Corynespora cassiicola*

**DOI:** 10.1101/2022.05.17.492372

**Authors:** Thaís Carolina da Silva Dal’Sasso, Vinícius Delgado da Rocha, Hugo Vianna Silva Rody, Maximiller Dal-Bianco Lamas Costa, Luiz Orlando de Oliveira

**Author notes:** Corresponding author: Luiz Orlando de Oliveira.

## Abstract

Effectors are secreted by plant-associated microorganisms in order to modify the host cell physiology. As effectors, the Necrosis- and Ethylene-inducing peptide 1-like proteins (NLPs) are involded in the early phases of plant infection and may trigger host immune responses. *Corynespora cassiicola* is a polyphagous plant-pathogen that causes target spot on many agriculturally important crops. Using genome assembly, gene prediction, and proteome annotation tools, we retrieved 135 NLP-encoding genes from proteomes of 44 isolates. We explored the evolutionary history of NLPs using Bayesian phylogeny, gene genealogies, and selection analyses. We accessed the expression profiles of the NLP genes during the early phase of *C. cassiicola*–soybean interaction. Three NLP effector genes (Cc_NLP1.1, Cc_NLP1.2A, and Cc_NLP1.2B) were maintained in the genomes of all isolates tested. A non-effector NLP gene (Cc_NLP1.3) was found in three isolates that had been originally obtained from soybean. NLP effectors were under different selective constraints: Cc_NLP1.1 was under stronger selective pressure, while Cc_NLP1.2A was under a more relaxed constraint. Meanwhile, Cc_NLP1.2B likely evolved under either positive or balancing selection. Despite highly divergent, the effector NLPs maintain conserved the residues necessary to trigger plant immune responses, suggesting they are potentially functional. Only the Cc_NLP1.1 effector gene was significantly expressed at the early hours of soybean colonization, while Cc_NLP1.2A and Cc_NLP1.2B showed much lower levels of gene expression.

## INTRODUCTION

Effectors are molecules secreted by plant-associated pathogens in order to modify the physiology of the host cell; by doing so, they facilitate the infection process (Lo Presti et al. 2015). The adaptation to new hosts and the evasion of recognition require the pathogen to acquire continuous innovations, such as changing the effector repertoires and recruiting rapidly evolving genes (Menardo et al. 2017; Fouché et al. 2018; Sánchez-Vallet et al. 2018). The discovery of effector proteins that share certain conserved domains provided new investigative tools for the study of plant-pathogen interactions. The *in silico* characterization of domains makes possible to unravel putative effectors based on genome annotations; therefore, allowing for the prediction of the likely function of the pathogen proteins in the host cell (Sonah et al. 2016; Jones et al. 2018). The presence of conserved regions generally defines proteins belonging to a gene family; those conserved regions can be share even by phylogenetically distant organisms (Franceschetti et al. 2017; Liu et al. 2019).

The Necrosis- and Ethylene-inducing peptide 1-like proteins (NLPs) constitute a superfamily of proteins present in a wide range of plant-associated microorganisms, including fungi, oomycetes, and bacteria (Gijzen and Nürnberger 2006; Oome and Van Den Ackerveken 2014; Seidl and Van den Ackerveken 2019). The NLPs are usually considered effectors owing to their cytotoxicity to eudicot plants that induces both plant immune responses and cell death by necrosis (Fellbrich et al. 2002; Bae et al. 2006; Qutob et al. 2006; Lenarčič et al. 2017; Ono et al. 2020). Although noncytolytic NLPs cannot permeabilize the plant membrane, they retain the capability of triggering plant immune responses (Qutob et al. 2006; Ottmann et al. 2009; Cabral et al. 2012; Dong et al. 2012; Böhm et al. 2014; Oome et al. 2014).

The presence of an NPP1 domain (Pfam PF05630) characterizes the NLPs (Fellbrich et al. 2002). Within its central part, the NPP1 domain display a highly conserved amino acid motif (GHRHDWE), which is involved in cation binding (Gijzen and Nürnberger 2006; Ottmann et al. 2009; Böhm et al. 2014; Oome et al. 2014). Based on the number of conserved cysteines in the protein, NLPs have been classified in three types: Type I, II, and III, respectively (Gijzen and Nürnberger 2006; Oome and Van Den Ackerveken 2014). Type I NLPs is the most frequent type; members of this type have two conserved cysteines that are important for the formation of a disulfide bridge (Gijzen and Nürnberger 2006). A variant called Type Ia differs from Type I by the occurrence of distinct amino acid substitutions and have been documented among some oomycetes (Oome and Van Den Ackerveken 2014). Type II NLPs possess four cysteines and formed two disulfide bridges (Gijzen and Nürnberger 2006; Oome and Van Den Ackerveken 2014). Type III NLPs usually display six cysteines residues and has been described in some ascomycetes and bacteria (Oome and Van Den Ackerveken 2014; Seidl and Van den Ackerveken 2019). The broad distribution of the three NLP types in the Ascomycota raised the possibility that the superfamily may have originated within this phylum (Oome and Van Den Ackerveken 2014).

Previously, we investigated the evolutionary history of NLPs across the Dothideomycetes, the largest class of ascomycetes (Dal’Sasso et al. 2022). We found evidence for a phylogeny-based classification of the NLP types into distinct NLP families: NLP1, NLP2, and NLP3. In addition to the consistently small size of the NLP superfamily among Dothideomycetes, we found that the NLP1 family was the most predominant family across a variety of taxa and could be split further into two subfamilies: NLP1.1 and NLP1.2 (Dal’Sasso et al. 2022). Amongst 79 species of Dothideomycetes, *Corynespora cassiicola* was one of the few species in which the members of both NLP1.1 and NLP1.2 subfamilies were present simultaneously. Namely, the genome of *C. cassiicola* possesses one gene that is member of the NLP1.1 subfamily and two genes that are members of the NLP1.2 subfamily (Dal’Sasso et al. 2022).

*Corynespora cassiicola* (Berk. & M. A. Curtis) C. T. Wei is usually referred as a necrotrophic pathogen due the necrotic lesions it causes on plants (Lopez et al. 2018). However, some isolates can be taken as endophytes and others as saprophytes (Déon et al. 2012b, 2014; Lopez et al. 2018). *Corynespora cassiicola* is one of the most destructive pathogens of *Hevea brasiliensis* (rubber tree) in East Asia; causing the *Corynespora* Leaf Fall (Déon et al. 2012a; Lopez et al. 2018). On many host species, this disease is referred to as ‘target spot’ because the pathogen causes necrotic circular lesions with a cracked center that resemble a target (Dixon et al. 2009; Sumabat et al. 2018). Target spot induces premature defoliation in a number of crops, which in turn reduces yield and results in economic losses. *Glycine max* (soybean), *Gossypium hirsutum* (cotton), *Cucumis sativus* (cucumber), and *Carica papaya* (papaya) are a few of the plant hosts to which *C. cassiicola* causes economically important diseases (Dixon et al. 2009; Déon et al. 2014; Sumabat et al. 2018; Gao et al. 2020; Dal’Sasso et al. 2021). In rare cases, *C. cassiicola* has been reported associated with human infection (Looi et al. 2017).

*Corynespora cassiicola* may comprise several phylogenetic lineages, each of which presenting different degrees of association with host specialization, pathogenicity and growth rate, but not with the geographical origin of the isolates (Dixon et al. 2009; Déon et al. 2014). Levels of host specialization suggest the involvement of specialized effectors that are recognized by a limited range of compatible hosts (Lopez et al. 2018; Gao et al. 2020). So far, the necrotrophic toxin cassiicolin is the only completely characterized effector of *C. cassiicola* (Breton et al. 2000; de Lamotte et al. 2007; Déon et al. 2014; Lopez et al. 2018; Ngo et al. 2022). Cassiicolin is an accessory effector, which means it is lineage specific and may not occur in all *C. cassiicola* isolates (Déon et al. 2014; Lopez et al. 2018; Wu et al. 2018). In addition to the cassiicolin-encoding genes, approximately 90 other effector genes are differentially expressed during *C. cassiicola* infection of rubber leaves; an NLP-encoding gene is present among them (Lopez et al. 2018). The role of NLP as effectors and whether all members of the NLP family in *C. cassiicola* are involved during infection remain elusive.

Herein, we investigated the evolutionary history of NLPs within a species framework; we used the plant pathogen *C. cassiicola* as model organism. First, we used standard genome assembly, gene prediction, and proteome annotation to retrieve NLP-encoding genes from 44 isolates of *C. cassiicola* that were available in public repositories. Next, we explored the genealogy of NLP-encoding genes using Bayesian phylogenetic and gene genealogy tools. Subsequently, we used statistical tests to investigate how selection shaped the NLP effector-encoding genes of *C. cassiicola*. Finally, we accessed the expression profiles of the NLP genes during the early phase of *C. cassiicola*–soybean interaction. Our investigation allowed us to address the following four questions: a) To what extent is the NLP family size conserved among isolates of *C. cassiicola*? b) Does each member of the NLP gene family encode for a putative effector protein? c) What evolutionary constraints shaped the evolution of NLP genes in *C. cassiicola*? d) What expression patterns the NLP genes of *C. cassiicola* exhibit during the early phase of soybean colonization?

Unveiling processes that drive the evolution and diversification of NLP-encoding genes in *C. cassiicola* will provide crucial insights into the understanding of the evolution of effector genes; therefore, shedding light into how NLPs may contribute to the pathogenicity profiles of this ubiquitous plant pathogen.

## MATERIAL AND METHODS

### Data assembly

We assembled both protein and coding-DNA sequence (CDS) data from 44 isolates of *C. cassiicola*. From GenBank, we downloaded genome sequences from 42 fungal isolates and predicted their proteins and associated CDSs. We also included protein sequences and CDSs from additional two isolates: CCP (Lopez et al. 2018) and CC_29 (Dal’Sasso et al. 2021). Thus, our set contained 44 isolates of *C. cassiicola*; hereafter, this set was referred to as “the Corca set” (Table S1).

### *De novo* genome assembly and gene prediction

Genomic data from two isolates of *C. cassiicola* (India_Hevea and TScotton1) were available on GenBank as raw reads only; therefore, they required *de novo* genome assembly prior to subsequent analyses. We evaluated the quality of the raw reads using FastQC v0.11.5 (https://www.bioinformatics.babraham.ac.uk/projects/fastqc/). Prior to genome assembly, we ran Trimmomatic v0.36 (Bolger et al. 2014) using the settings “ILLUMINACLIP:TruSeq3.fa:2:30:10”, “LEADING:3”, “TRAILING:3”, and “MINLEN:50” to remove adaptors and low quality (Q < 30) reads. *De novo* assembly was carried out using SPAdes v3.9.0 (Bankevich et al. 2012) for Illumina reads, with the parameters set as “-k 21,33,55,77” for isolate India_Hevea and “-k 21,33,55” for isolate TScotton1, and the parameter “--careful” for both isolates. We accessed the quality of genome assemblies using QUAST v5.0.2 (Mikheenko et al. 2018), with default parameters for eukaryotes. Finally, we assessed the completeness of core fungal orthologs using BUSCO v4.0.5 (Simão et al. 2015) based on the dataset fungi_odb10 with 758 core orthologs.

For those genomes we had downloaded previously from GenBank, gene prediction was peformed using Augustus v3.2.2 (Stanke and Morgenstern 2005). For training Augustus, we used the pre-existing gene structures available for the isolate CCP. Training was conducted according to manual instructions. Subsequently, we ran Augustus using isolate CCP as ‘reference species’ with parameters “-- codingseq”, “--protein”, and “--cds”.

### Protein annotation and assemble of NLP homologues

Protein annotation and assemble of NLP homologues were performed as previously described for Dothideomycetes (Dal’Sasso et al. 2022). The predicted proteomes of the Corca set was annotated using PfamScan with Pfam v32.0 (El-Gebali et al. 2019) and InterProScan v5.30.69 (Jones et al. 2014) with the following eight parameters: SMART-7.1, SUPERFAMILY-1.75, ProDom-2006.1, CDD-3.16, TIGRFAM-15.0, Pfam v31.0, Coils-2.2.1, and Gene3D-4.2.0.

Secreted proteins were defined by the presence of a signal peptide (according to SignalP v4.1; Petersen et al. 2011) and the absence of transmembrane domains (according to TMHMM v2.0; Krogh et al. 2001). Subsequently, TargetP (Emanuelsson et al. 2007) predicted the subcellular localization of the predicted secretome.

From the predicted proteomes, HMMER v3.2.1 (http://hmmer.org) predicted the NLP homologues (E-value < 0.001) using profile hidden markov models (HMMs) for NPP1 domain (PF05630). We declared a protein to be an NLP when the NPP1 domain was annotated by at least two out three softwares: PfamScan, InterproScan, and HMMER. We considered a given NLP to be an ‘effector NLP’ when it passes the following three tests: (a) SignalP predicted it harbored a signal peptide, (b) TMHMM predicted it had no transmembrane domain, and (c) TargetP predicted it was part of a secretoy pathway. When a given NLP failed any of these three tests, we regarded it to be a ‘non-effector NLP’. At this point, our algorithm sorted NLPs into either effector NLPs or non-effector NLPs.

Finally, we subject the predicted NLPs to EffectorP v3.0, which uses machine learning in order to classify proteins into three classes (effectors, unlikely effectors, or non-effectors) and to predict the localization of the effectors into the host cell (cytoplasm or apoplast) (Sperschneider and Dodds 2022).

### Orthogroups and Bayesian phylogenies of NLPs

OrthoFinder v2.3.3 (Emms and Kelly 2015) established orthologous relationships among the 44 members of the Corca set (Table S1) and calculated length and phylogenetic distance-normalized scores from an all-versus-all alignment using DIAMOND v0.9.24 (Buchfink et al. 2014). OrthoFinder assumed ortholog groups (orthogroups) by Markov clustering analysis.

We assembled an additional dataset, which contained the predicted NLPs (CDSs) that were present in the Corca set. This dataset of predicted NLPs was referred to as “the NLP set”. Bayesian phylogenetic analyses based on the NLP set allowed for the reconstruction of the phylogenetic relationships among the NLPs of *C. cassiicola*.

Alignments were obtained using the L-INS-I method, as implemented in MAFFT v7.453 (Katoh and Standley 2013). During alignment of CDSs, we detected the presence of an insertion of either 105 bp (in isolate CC_29) or 153 bp (in isolates 777AA and RUD). In order to find out their origin, these insertions were used as query in local BLASTn (E-value < 1e-4) to search for homologous sequences within the Corca set and within the repeat library we have built previously for isolate CC_29 (Dal’Sasso et al. 2021).

According to BIC, jModelTest v2.4 (Darriba et al. 2012) indicated GTR+I as the best fit model for the NLP set. Subsequently, phylogenetic analyses were conducted on MrBayes v3.1.2 (Ronquist and Huelsenbeck 2003), using two simultaneous runs, one cold chain and seven heated chains in each run. The number of generations was five million and the temperature was 0.25. Trees were sampled every 5000 generations and the first 250 trees were discarded as burn-in samples. Convergence was diagnosed as follows: a) the standard deviation of split frequencies at the end of each run (below 1.5%; according to the output from MrBayes) and b) the Effective Sample Size (ESS) of each parameter (above 200 for all statistics) in Tracer v1.7.1 (Rambaut et al. 2018). For each analysis, a 50% majority-rule consensus tree of the two independent runs was obtained with posterior probabilities (PP) that were equal to bipartition frequencies. The phylogenetic tree was visualized in FigTree v1.4.4 (http://tree.bio.ed.ac.uk/software/figtree/).

### Network analysis and measures of nucleotide diversity

Gene genealogies of predicted NLP encoding-genes were inferred using the median-joining network method (Bandelt et al. 1999), as implemented in Network v4.613 (Fluxus Technology Ltd.) with default parameters. Preliminary analyses indicated that sequence alignments among introns were difficult to achieve; therefore, we chose to carry out the network analyses using CDS data only.

From the NLP set, we inferred the genealogical relationships among the full set of predicted NLP genes (CDSs). Afterwards, we made partitions within the NLP set according to the sub-clades we just recovered during the preceding phylogenetic analysis. There were four sub-clades, which were referred to as Cc_NLP1.1, Cc_NLP1.2A, Cc_NLP1.2B, and Cc_NLP1.3, respectively. Each sub-clade underwent an additional round of haplotype network analysis. Finally, we used DNAsp v6 (Rozas et al. 2017) to estimate the following five measures of molecular diversity within each sub-clade: H, number of haplotypes; H_d_, haplotype diversity; π, nucleotide diversity; K, average number of nucleotide differences; S, number of polymorphic sites.

### Selection analyses

We applied five different statistical tests to investigate how selection shaped the NLP effector-encoding genes of *C. cassiicola*. The neutrality tests of Tajima’s D (Tajima 1989), Fu and Li’s D* and F* (Fu and Li 1993) were carried out in DNAsp v6. They tested for departures from neutrality based on allele frequency distribution indices (Ramírez-Soriano et al. 2008). As input for the analyses, we used the partition of the NLP set. For those analyses, we removed the nucleotides responsible to encode the signal peptides.

The McDonald and Kreitman test (MKT) (McDonald and Kreitman 1991) tested the hypothesis of positive selection in the predicted NLP effector genes of *C. cassiicola*. The MKT calculated the ratio of the number of non-synonymous polymorphic sites (*Pn*) by the number of synonyms polymorphic sites (*Ps*) within the species and compared to the ratio of the number of non-synonymous nucleotide substitutions (*Dn*) by the number of synonymous nucleotide substitutions (*Ds*) between species; therefore, an outgroup was required to determine in which sites the differences were fixed (Bhatt et al. 2010; Rody and Oliveira 2018). The MKT also calculated the Neutrality Index (NI), which indicates how far polymorphism is from neutral evolution, and the rate of positive selection (α), which positive values corroborates with positive selection. The MKT analyses were carried out using the partitions we had obtained previously. To choose the outgroup we returned to the results of the phylogenetic analyses of NLPs from Dothideomycetes we had performed previously (Dal’Sasso et al. 2022) and selected as outgroup the closest orthologue protein to each of the NLPs of *C. cassiicola*. As outgroups, we added to our datasets the following homologous sequences: Stano_3364 (from *Stagonospora nodorum*) homologous to Cc_NLP1.1, Bimnz_484389 (*Bimuria nova-zelandiae*) to Cc_NLP1.2A, and Cocsa_135644 (*Cochliobolus sativus*) to Cc_NLP1.2B. For those analyses, we removed the nucleotides that would give rise to the signal peptides.

The package *codeml* from PAML v4.9j (Yang 2007) carried out tests of positively selected sites in the effector NLPs. The package calculates the ratio of nonsynonymous (d*N*, amino acid changing) to synonymous (d*S*, amino acid retaining) substitution rates (ω = d*N*/d*S*) (Rody and Oliveira 2018). Sites that are subject to natural selection show dN/dS = 1, whereas those that are under purifying selection and those that are evolving by positive selection have dN/dS < 1 and dN/dS > 1, respectively (Terauchi and Yoshida 2010). To ensure that ω variation represented amino acid sites that were fixed along independent lineages, we used the same inputs as for MKT. Possible ambiguities and alignment gaps were cleanup by choosing option cleandata = 1. A phylogenetic tree for each NLP effector-encoding gene was generated by PhyML v3.1 (Guindon and Gascuel 2003), using the GTR model of nucleotide substitution, and used as input along with the aligned datasets. We performed likelihood ratio tests (LTRs) between two pairs of models: M2 (selection) against M1 (neutral) and M8 (beta&ω) against M7 (beta). Positively selected codon sites were predicted using the Bayes Empirical Bayes (BEB) method. Condon sites were considered under positive selection when ω > 1 and posterior probability calculated by the BEB was > 95%.

### Motif searching in the effector NLPs of *C. cassiicola*

The packages GLAM2SCAM and GLAM2, as implemented in the MEME suite v5.1.0 (Bailey et al. 2009), created a consensus logo of the effector NLPs in *C. cassiicola*. The package GLAM2 searched for pattern of motifs among the predicted effector NLPs; while GLAM2SCAM searched for matches for the motif initially discovered by GLAM2. Each match received a score, indicating how well it fit the motif. GLAM2 returned a logo with the consensus sequence with the highest score.

Visualization of the 3D structure of the effector NLP was obtained in PyMOL v2.3 (Schrödinger, LLC), using as input the PBD format file from *Pythium aphanidermatum* that was available under accession 3GNU at the Protein Data Bank. The protein sequence was depicted as a cartoon diagram and color coded. In the cartoon, the sites that *codeml*/PAML suggested to be under positive selection were highlighted.

### Expression of the NLP genes of *C. cassiicola*

Using RT-qPCR, we investigated the expression *in planta* of the predicted NLP genes of *C. cassiicola*. We used *C. cassiicola* isolate CC_29 (originally isolated from soybean leaves; Dal’Sasso et al. 2021) because it also contained a copy of the rare non-effector Cc_NLP1.3 in addition to the widespread Cc_NLP1.1, Cc_NLP1.2A, and Cc_NLP1.2B copies.

Plants of soybean cultivar “MG/BR-46 Conquista” were cultivated in a greenhouse facility until they reached the stage V3. Inoculation procedures were as follows: four droplets (20 μl each) of a conidia suspension (3.5 × 10^4^ conidia/ml) from isolate CC_29 amended with 0.01% Tween 20 were spotted on the abaxial face of each leaflet from trifoliate leaves. Immediately after the inoculation, plants were incubated in a humid chamber at 25 ± 2 °C. Controls consisted of plants that had leaflets inoculated with sterile distilled water supplemented with 0.01% Tween 20. Sampling consisted of the collection of four discs (1.4 cm^2^) per leaflet (at the inoculation spots) using a cork borer at 0, 6, 12, and 18 hours post inoculation (hpi). For each time point, three independent biological repeats were collected, each of which from a distinct plant. Samples were immediately frozen in liquid nitrogen and stored at -80°C until subsequent RNA extraction.

Total RNA was extracted from each sample using TRIzol reagent (Life Technologies) according to manufacturer’s specifications. RNA was quantified on a NanoDrop 2000 spectrophotometer (Thermo Fisher Scientific) and visualized on 1.5% (w/v) agarose gel. A total of 4 μg of RNA was treated with one unit of DNase I Amplification Grade (Invitrogen) to remove potential contamination with genomic DNA. The cDNA synthesis was performed using 4 μg of total RNA, Oligo(dT)_18_ primers, and the M-MLV Reverse Transcriptase (Invitrogen) following the manufacturer’s instructions.

Based on the genome of isolate CC_29 (Dal’Sasso et al. 2021) and using Primer Blast (https://www.ncbi.nlm.nih.gov/tools/primerblast/), we designed primers for each of the four predicted NLP genes (CDSs) and for the β-tubulin gene (CDS). We checked primer quality using NetPrimer (http://www.premierbiosoft.com/netprimer/). Primers used in this study are listed (Table S2).

Amplifications (RT-qPCR) were performed on a 7500 Real-Time PCR System (Applied Biosystems) using Power SYBR Green PCR Master Mix (Life Technologies), specific primers (Table S2), and 1 μl of a 2-fold diluted cDNA. The reactions were performed as follows: 2 min at 50 °C, 10 min at 95 °C, and 40 cycles of 94 °C for 15 seconds, and 60 °C for 1 min. To confirm quality and primer specificity, we visually inspected the T_m_ (melting temperature) of amplification products in dissociation curves for each primer individually.

For quantitation of NLP genes, we used β-tubulin as the endogenous control gene for data normalization. Gene expression was quantified using the 2^-ΔCt^ method for absolute quantification. To confirm the observed differences in gene expression, we applied a one-way ANOVA followed by Tukey’s test (*p*-value < 0.05) for each NLP gene, independently. Statistical analyses were implemented in R v3.6.2 (http://www.R-project.org/). A heatmap was designed using the *tidyr* and *ggplot2* packages, also in R.

## RESULTS

### *De novo* genome assembly

*De novo* genome assembly of isolates India_Hevea and TScotton1, available on GenBank as raw reads, resulted in good quality genomes (Table S3). The final genome assemblies of India_Hevea and TScotton1 added up to 42.89 Mbp (arranged in 1,059 scaffolds; ≥ 500 bp) and 42.11 Mbp (2,188 scaffolds; ≥ 500bp), respectively. The BUSCO analysis identified 99.6% of the orthologs as complete for India_Hevea, and 98.9% as complete for TScotton1. The BUSCO scores indicated a high level of completeness, making the genomes suitable for the downstream analyses.

### Phylogenetic relationships among NLP-encoding genes

The genome sequences of 44 isolates of *C. cassiicola* yielded a total of 135 predicted NLP-encoding genes (CDSs). The phylogenetic tree (Fig. 1) showed maximally supported nodes (posterior probabilities = 100, for all main nodes). The 135 copies were grouped into four major clades. Based on OrthoFinder analysis, each of the four major clades corresponded to an orthogroup.

**Fig. 1.**
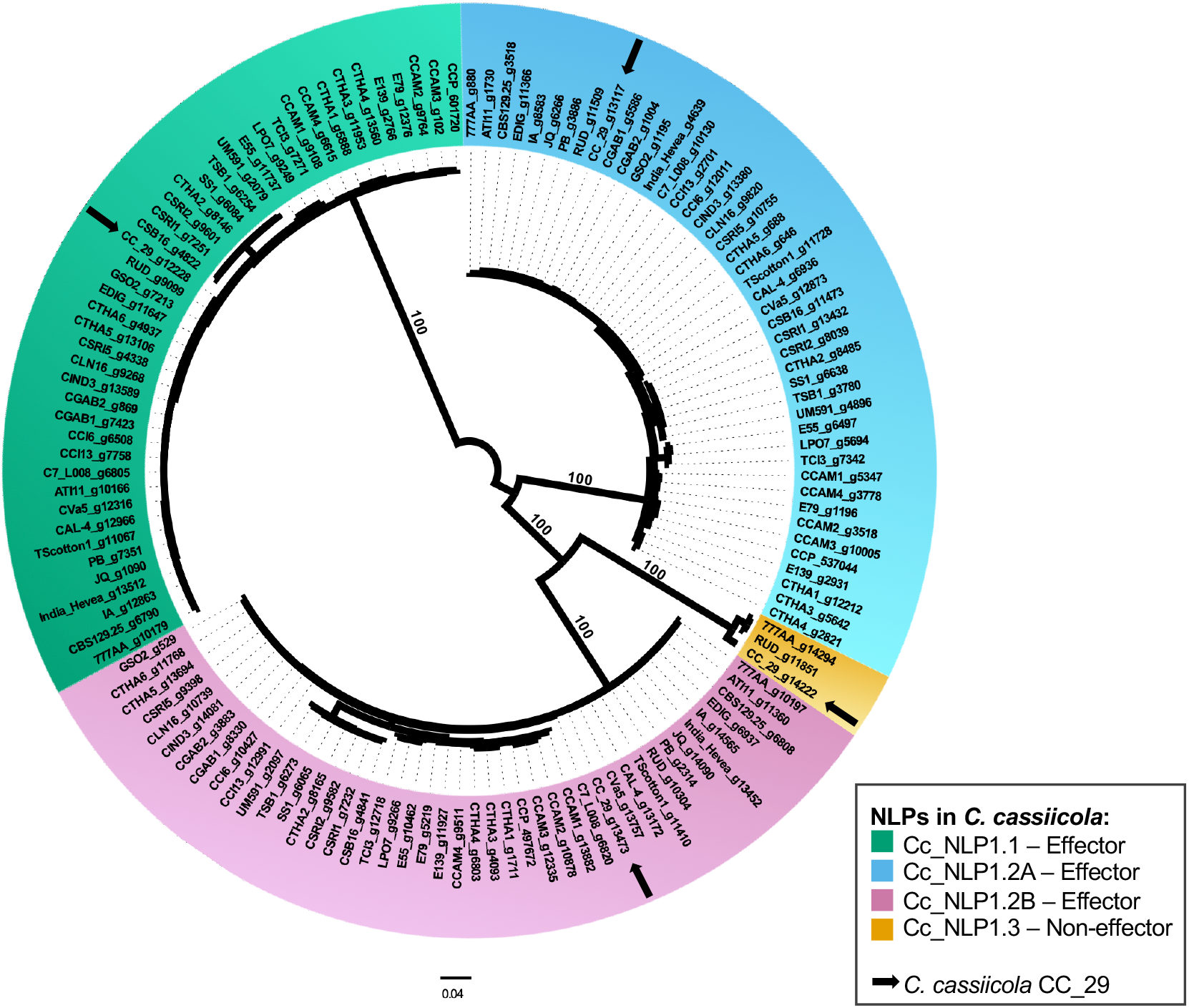
Phylogenetic relationships among 135 predicted Necrosis- and Ethylene-inducing peptide 1-like protein (NLP)-encoding genes (CDS) in *Corynespora cassiicola*. The DNA dataset for the unrooted Bayesian phylogeny (consensus tree) was 900 base pairs long and contained NLPs from 44 isolates. Arrows indicate the placements of the four predicted NLP-encoding genes of *C. cassiicola* isolate CC_29. Along the tree, gene ID according to gene prediction using Augustus (or MycoCosm nomenclature for isolate CCP) followed the name of the isolate and identified the predicted NLP-encoding genes. Gene names are color-coded according to the CDS homology to any of the three effector NLPs or to one non-effector NLP of *C. cassiicola* CC_29, as indicated. Branch lengths are drawn to scale; nodal support values are given as posterior probabilities (when ≥ 95%) above the branches. Scale bar corresponds to the expected number of substitutions per site.

Each of the three copy-richest clades shared homology to one of the three NLPs (CCP_601720, CCP_537044, and CCP_497672) that were present in the genome of the isolate CCP (Dal’Sasso et al., 2022). Therefore, we named each of those three major clades after the sub-clades (subfamilies) within the NLP1 family we had recovered previously (Dal’Sasso et al. 2022): Cc_NLP1.1, Cc_NLP1.2A, and Cc_NLP1.2B (Fig. 1). Each of the 44 isolates had one copy of NLP-encoding gene from each of the three major clades. Those 132 copies were predicted to be NLP effector-encoding genes, that is, they encode for effector NLPs (proteins predicted to harbor a signal peptide each, no transmembrane domain, and part of a secretoy pathway). According to EffectorP classification, those 132 copies encoded NLPs that were predicted to be apoplastic effectors.

In addition to the three widespread genes, we predicted a fourth gene in the genomes of three isolates obtained from soybean that was cultivated in Brazil: CC_29 (gene CC_29_g14222), 777AA (777AA_g14294), and RUD (RUD_g11851).

In the phylogenetic tree (Fig. 1), copies of the fourth gene clustered together to form the fourth major clade (hereafter referred to as Cc_NLP1.3), which exhibited a sister relationship to the clade Cc_NLP1.2B.

Copies within the Cc_NLP1.3 clade showed features that distinguish them apart from the copies within the other three major clades. In addition to harboring many exclusive synonymous and non-synonymous substitutions, they carried a large insertion at the 5’ region, just after the predicted start codon (105 bp in CC_29_g14222; 153 bp in both 777AA_g14294 and RUD_g11851). BLASTn searches using each of those insertion sequences as queries did not find significant matches in the genomes of the remaining 41 isolates of *C. cassiicola* (Corca set; Table S1). Moreover, additional BLASTn searches did not find evidence for homology between those insertions and the repeat/transposable elements that were present in the genome of isolate CC_29 (Dal’Sasso et al. 2021). Finally, copies within the Cc_NLP1.3 clade were predicted to be non-effector NLP-encoding genes.

### Gene genealogies among NLP-encoding genes

The full haplotype network recovered 39 haplotypes among the 135 sequences of NLP-encoding genes (CDSs) that had been retrieved from genomes of 44 isolates of *C. cassiicola* (Fig. 2). The network exhibited four major sets of haplotypes (hereafter, each major set is referred to as a ‘haplogroup’). Without exception, both the topology of the network and the composition of each haplogroup were congruent with the four clades we had obtained in the preceding phylogenetic analysis (Fig. 1). To be consistent within our previous findings, we named the four haplogroups after the four phylogenetic clades: Cc_NLP1.1, Cc_NLP1.2A, Cc_NLP1.2B, and Cc_NLP1.3. While Cc_NLP1.1 and Cc_NLP1.3 occupied each a tip position on opposed ends of the full network, Cc_NLP1.2A and Cc_NLP1.2B occupied its center. The largest number of mutational steps between nearest haplogroups occurred between Cc_NLP1.1 and Cc_NLP1.2A (314 steps); while the smallest number occurred between Cc_NLP1.2B and Cc_NLP1.3 (234 steps). Distances between nearest haplogroups far exceeded the distance between nearest haplotypes within the haplogroups. Therefore, we split the full dataset according to haplogroups and analyzed each haplogroup independently.

**Fig. 2.**
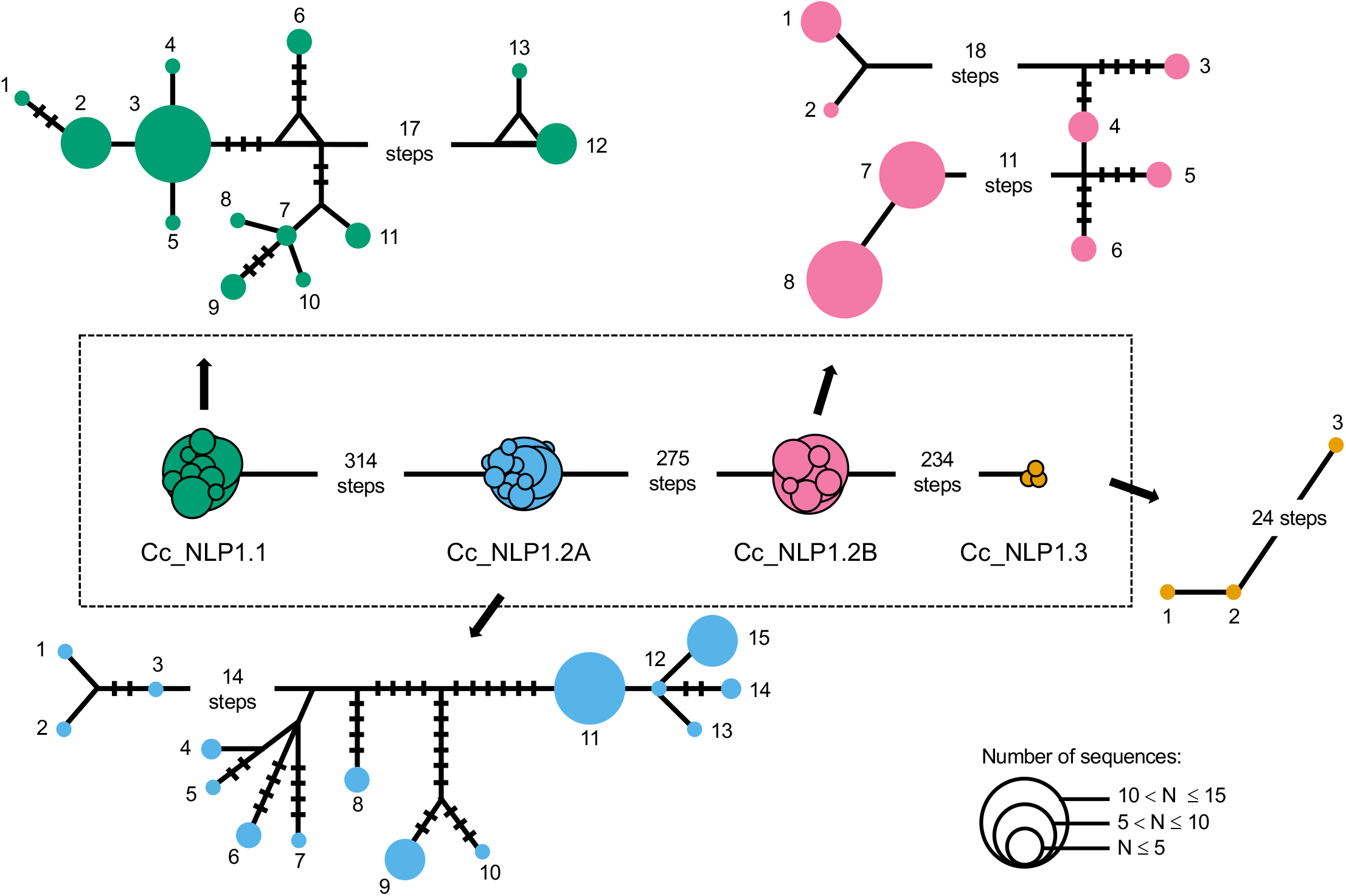
Genealogical relationships of the four haplogroups of predicted Necrosis- and Ethylene-inducing peptide 1-like protein (NLP)-encoding genes (CDS) of *Corynespora cassiicola*. Median-joining networks of each of the four haplogroups are color-coded according to the predicted NLP encoding-genes: green, Cc_NLP1.1; blue, Cc_NLP1.2A; pink, Cc_NLP1.2B; and orange, Cc_NLP1.3. A central box depicts the full set of haplogroups; the number of mutational steps is as indicated. A circle represents a given haplotype (coded with numbers); circle size is proportional to the relative frequencies, as indicated. Numbers of mutational steps are indicated with bars when more than one (unless indicated otherwise).

The four partial networks shed further light on the genealogical relationships among haplotypes within each of the haplogroups. Within each haplogroup, the mutational distances that separated two nearest haplotypes were short (up to 11, within Cc_NLP1.2B; up to 24 within Cc_NLP1.3). Mutational distances larger than three arise owing to missing intermediates. Those missing intermediates represent either extinct haplotypes or, more likely, extant haplotypes that may exist in the population of isolates yet to be sampled. Within each haplogroup, most of the missing intermediates occurred in tandem. The presence of large number of autapomorphic substitutions suggested that most of the NLP-encoding genes have experienced relatively high levels of genetic differentiation since their origin.

Measures of nucleotide diversity also indicated that the four haplogroups of NLP-encoding genes of *C. cassiicola* followed relatively distinct evolutionary paths (Table 1). Genetic differentiation reached the lowest values among members of haplogroup Cc_NLP1.1, with only ∼9 mutations as the average number of nucleotide differences (K) among pair of haplotypes. Additionally, nucleotide diversity (π) was also lower among members of Cc_NLP1.1. On the opposite end, genetic differentiation reached the highest values among members of haplogroup Cc_NLP1.3, in which pairs of haplotypes diverged by about 17 mutations on average, showing substantially large values of haplotype (H_d_) and nucleotide diversity (π).

**Table 1.** Measures of molecular diversity for the four haplogroups of NLP encoding-genes (CDS) in *Corynespora cassiicola*.

| NLP Haplogroup | # Sequences | # Variable Sites (S) | # Haplotypes (H) | Haplotype Diversity ($H_d$) | Nucleotide Diversity ($\pi$) | Average # of Nucleotide Differences (K) |
| --- | --- | --- | --- | --- | --- | --- |
| Cc_NLP1.1 | 44 | 35 | 13 | 0.86 | $1.3 \times 10^{-2}$ | 8.97 |
| Cc_NLP1.2A | 44 | 57 | 15 | 0.88 | $1.6 \times 10^{-2}$ | 11.68 |
| Cc_NLP1.2B | 44 | 39 | 8 | 0.83 | $1.6 \times 10^{-2}$ | 12.01 |
| Cc_NLP1.3 | 3 | 25 | 3 | 1.00 | $2.2 \times 10^{-2}$ | 16.67 |

### Selection analyses on NLP effector-encoding genes in *C. cassiicola*

We applied three selective neutrality tests (Tajima’s D, Fu and Li’s D*, and F*) on three sets of NLP effector-encoding genes (Cc_NLP1.1, Cc_NLP1.2A, and Cc_NLP1.2B). During those three tests, we used information from the CDSs only (Table 2, Fig. S1). Each selective test was carried out using both the full extension of the genes and a contiguous sliding window of 25 bases. The Tajima’s D test recovered non-significant values for all sets (*p*-value > 0.10), both for the full sequence extension and the contiguous sliding window. Thus, results of the Tajima’s D test supported the null hypothesis of neutrally evolving genes.

**Table 2.** Results of neutrality tests for three haplogroups of NLP effector genes (CDS) in *Corynespora cassiicola*.

| Effector Haplogroup | Tajima's | Fu and Li's | | McDonald and Kreitman | | | Positive selected amino acid sites ($\omega > 1$) (amino acid)<sup>2</sup> |
| --- | --- | --- | --- | --- | --- | --- | --- |
| | D (*p*-value) | D*<sup>1</sup> (*p*-value) | F*<sup>1</sup> (*p*-value) | NI | $\alpha$ | *p*-value | |
| **Cc_NLP1.1** | 0.01 (*p* > 0.10) | 0.79 (*p* > 0.10) | 0.62 (*p* > 0.10) | 0.16 | 0.84 | *p* < 0.01 | - |
| **Cc_NLP1.2A** | -0.47 (*p* > 0.10) | 0.26 (*p* > 0.10) | 0.01 (*p* > 0.10) | 0.17 | 0.83 | *p* < 0.001 | - |
| **Cc_NLP1.2B** | 1.15 (*p* > 0.10) | 1.63 (*p* < 0.02) | 1.73 (*p* < 0.05) | 0.52 | 0.48 | *p* > 0.10 | 36 (R)*, 77 (N)** |

<sup>1</sup> Asterisks indicate no outgroup
<sup>2</sup> The *codeml* tool performed predictions with model M8, as implemented in the PAML 4.9j software package. BEB calculated the Bayesian PPs (>95%\* or >99%\*\*).

The Fu and Li’s D* and F* tests provided similar results. The F* statistics are shown in Fig. S1. Using the full extension of the genes, Fu and Li’s D* and F* recovered non-significant values for two sets (Cc_NLP1.1 and Cc_NLP1.2A), with *p*-values > 0.10; and significant positive values for Cc_NLP1.2B (*p*-value < 0.02 for D*; *p*-value < 0.05 for F*) (Table 2). Therefore, the D* and F* tests rejected the null hypothesis of neutrally evolving genes and suggests that balancing selection is acting upon Cc_NLP1.2B. Using the sliding window approach, Fu and Li’s D* and F* recovered significant negative values for some nucleotides between the midpoints 616-650 of Cc_NLP1.2A (*p*-value < 0.10 for D*; *p*-value < 0.05 for F*), which indicates an excess number of alleles (Fig. S1).

Results of the MKTs showed significant values for Cc_NLP1.1 (*p*-value < 0.01) and Cc_NLP1.2A (*p*-value < 0.001), according to Fisher’s exact test and G-test, as implemented in DNAsp. However, results showed NI < 1 and α values 0 < α < 1 (Table 2) for all NLP effector genes (Cc_NLP1.1, Cc_NLP1.2A, and Cc_NLP1.2B). Taking together, MKT results indicate an excess of fixation of non-neutral substitutions on NLP effector-encoding genes and suggest that, to some extent, they evolved under positive selection.

To test for selective pressure upon amino acid sites, we evaluate positive selection using site-to-site models (M1 vs. M2a, M7 vs. M8), as implemented in the *codeml* tool from PAML software package. Congruent results were obtained for models M2 and M8 (both for selection). The results of the M8 model are shown (Table 2). In the Cc_NLP1.2B, two sites were identified to be under positive selection (PP > 95%) as calculated by the BEB approach (Table 2, Fig. 3). Positively selected sites referred to residue 36, which was an arginine (R36), and residue 77, which was an asparagine (N77). Both residues are located on regions that are predicted to be α-helices, located at one side of the protein and adjacent to the negatively charged (cation binding) cavity that forms around the conserved GHRHDWE motif (Fig. 3). The alternative allele for the positive selected residue 36 detained a lysine (K36), instead of an arginine (R36). For the positive selected residue 77, alternative alleles detained either a glycine (G77) or a serine (S77), instead of an asparagine (N77).

**Fig. 3.**
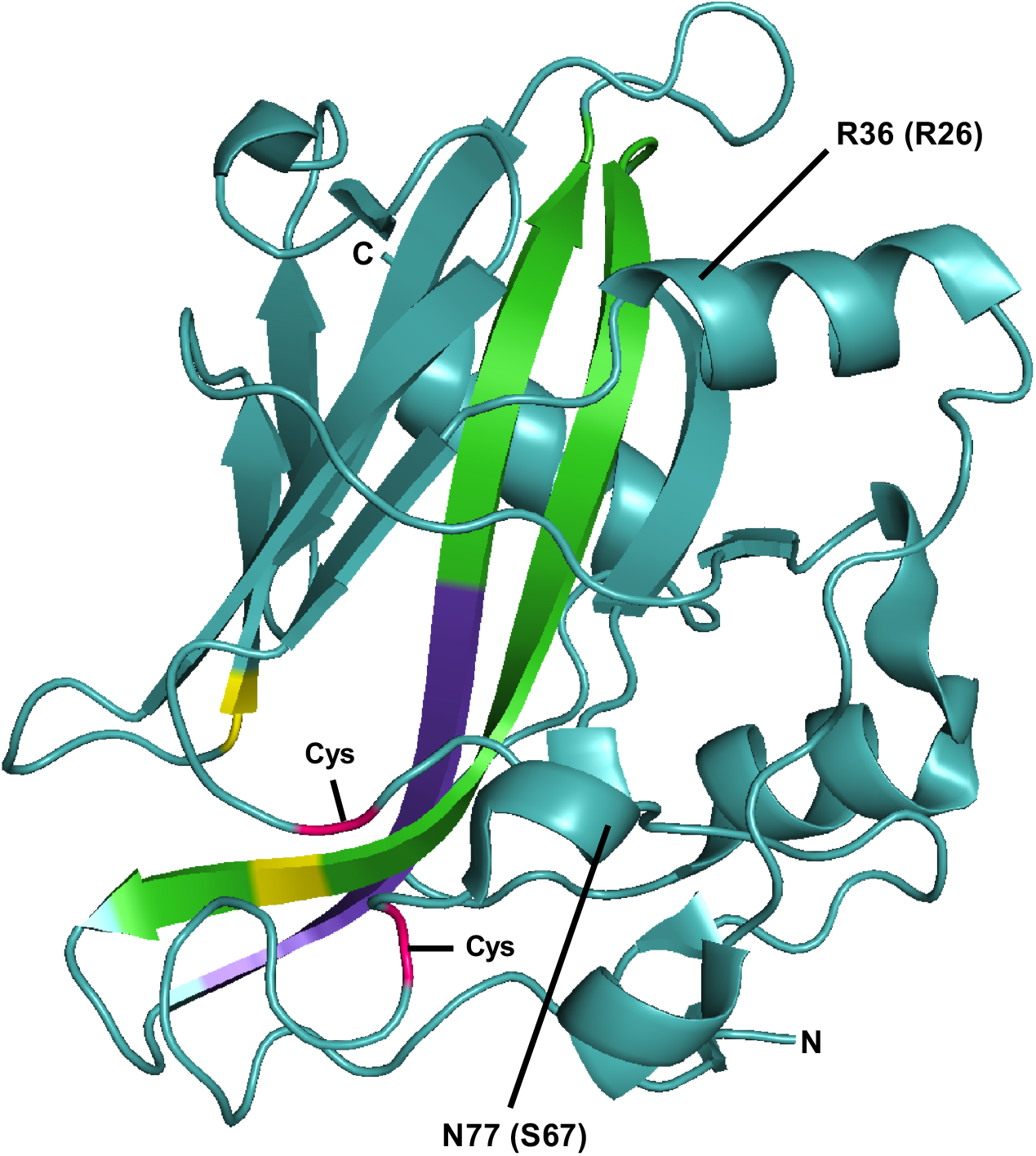
PyMOL diagrams depicting the 3D structure of the Necrosis- and Ethylene-inducing peptide 1-like (NLP) effector of *Pythium aphanidermatum* and the relative locations of two positively selected sites of the Cc_NLP1.2B effector in *Corynespora cassiicola*. Protein structure was taken from the Protein Data Bank (accession number 3GNU). According to *codeml*/PAML, there were two positively selected amino acid sites (R36 and N77) in Cc_NLP1.2B. R26 and N67 are the corresponding sites in the structure of the effector NLP of *Pythium aphanidermatum*. Color codes: purple, conserved GHRHDWE site; green, semiconserved sites in the central β-strands; yellow, residues directly involved in cation binding; pink; conserved cysteine residues involved in the protein stabilization (disulfide bridge formation). Letter codes: Cys, conserved cysteines residues; N, N-terminus; and C, C-terminus.

### Nucleotide substitutions within NLP effector-encoding genes

We documented the nature and context of nucleotide substitutions along the haplotypes within haplogroups Cc_NLP1.1, Cc_NLP1.2A, and Cc_NLP1.2B (Fig. S2). During each independent analysis, gene members we had identified in isolate CC_29 were taken as the reference sequences for that haplogroup: CC_29_g12228 (Cc_NLP1.1), CC_29_g13117 (Cc_NLP1.2A), and CC_29_g13473 (Cc_NLP1.2B).

Haplogroup Cc_NLP1.1 contained a total of 35 nucleotide substitutions among 13 haplotypes (Fig. S2A, Table 1). Among them, four (10%) were non-synonymous mutations, that is, they resulted in amino acid replacements. One of them was a conservative amino acid replacement at the C-terminal half of the NPP1 domain of haplotypes 11-13. The remaining three non-synonymous mutations were located outside the NPP1 domain, and corresponded to one conservative and two non-conservative amino acid replacements, respectively.

Haplogroup Cc_NLP1.2A exhibited the highest number of nucleotide substitutions; there were 57 of them among 15 haplotypes (Fig. S2B, Table 1). Amongst them, eight (14%) were non-synonymous mutations, which resulted in seven distinct amino acid replacements in the protein sequence. Two consecutive non-synonymous mutations resulted in a non-conservative amino acid replacement at the N-terminal half of the NPP1 domain, adjacent to the GHRHDWE motif. The remaining non-synonymous mutations corresponded to conservative amino acid replacements: five of them in the C-terminal half of the NPP1 domain and one immediately after the domain.

Among the eight haplotypes of haplogroup Cc_NLP1.2B, there were a total of 39 nucleotide substitutions (Fig. S2C, Table 1). Amongst the substitutions, eight (20%) were non-synonymous mutations and resulted in seven different amino acid replacements in the protein sequence. At the N-terminal half of NPP1 domain, a non-synonymous mutation resulted in conservative amino acid replacement (haplotypes 1-4). Also, in the N-terminal half of the NPP1 domain, two consecutive non-synonymous mutations resulted in three different amino acids replacements at the same codon (haplotypes 1-3, 5). According to the *codeml*/PAML analysis, the sites that harbor those amino acid replacements (either conservative or non-conservative mutations) corresponded to positively selected sites (Fig. 3, Table 2). The remaining five non-synonymous mutations corresponded to conservative amino acid replacements: one took part in the CDS portion encoding for the signal peptide, followed by one in the portion encoding between the signal peptide and the NPP1 domain. Finally, three conservative amino acid replacements took part in the CDS portion encoding for the NPP1 domain (one in the N-terminal half and two in the C-terminal half of the domain).

### Expression patterns of NLP-encoding genes

Using soybean as a host species, we used RT-qPCR analyses to characterize the expression patterns of the four NLP-encoding genes we found in the genome of the isolate CC_29. During the first 18 hpi, the NLP effector genes CC_29_g12228 (Cc_NLP1.1), CC_29_g13117 (Cc_NLP1.2A), and CC_29_g13473 (Cc_NLP1.2B) showed varying levels of expression (Fig. 4 and S3). Expression of the effector gene CC_29_g12228 (Cc_NLP1.1) took place very early, exhibited a significant increase between 12 and 18 hpi, and reached the highest level of absolute expression (fold variation) among the four members of the NLP family.

**Fig. 4.**
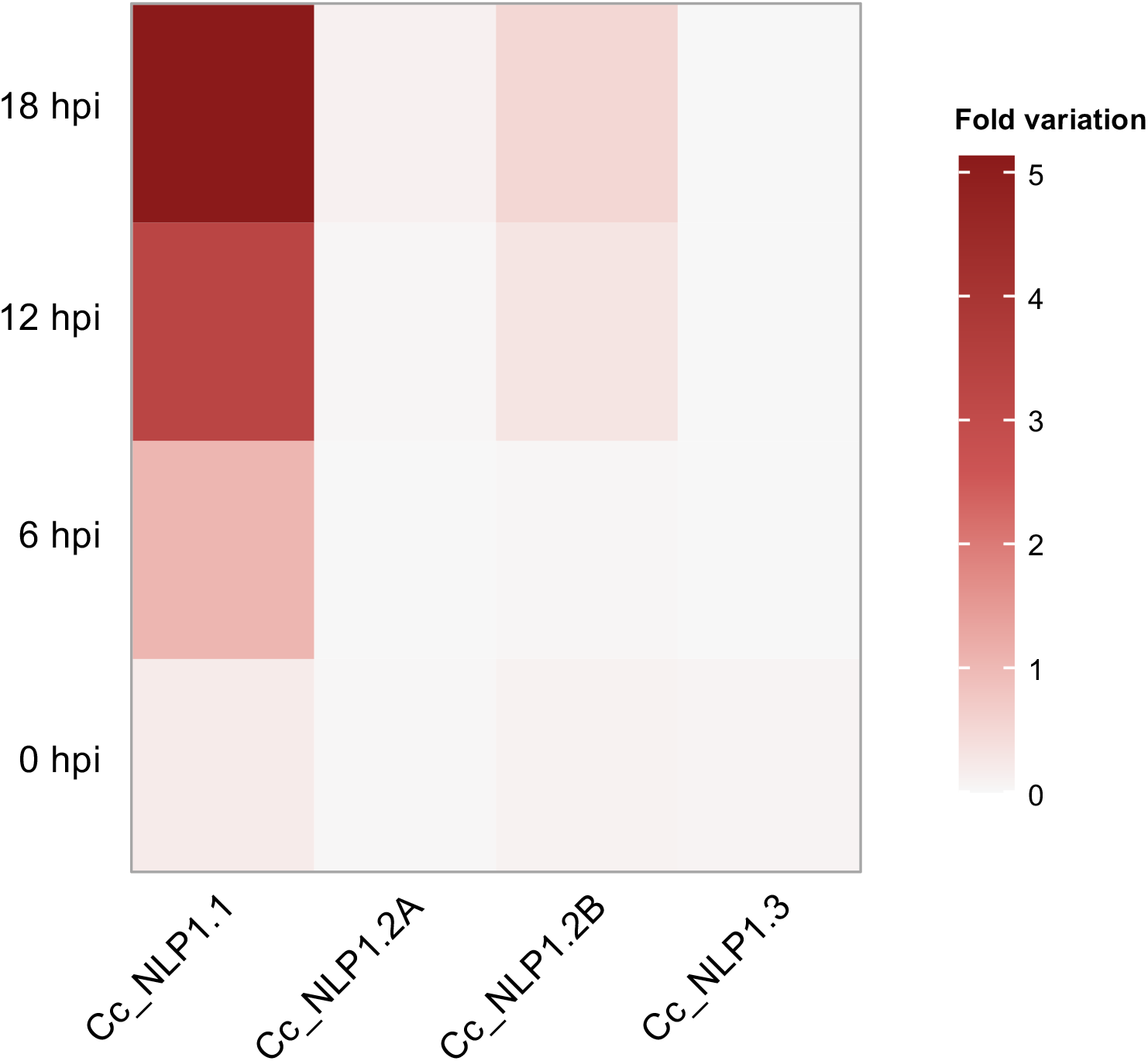
Expression patterns of the Necrosis- and Ethylene-inducing peptide 1-like (NLP) genes of *Corynespora cassiicola* (isolate CC_29) after inoculation in soybean plants. Gene expression was inferred by RT-qPCR analyses. The fold variation of gene expression of each NLP gene was calculated using the 2^-ΔCt^ method, the values were normalized to an endogenous control gene (β-tubulin). Columns correspond to NLP genes (from left to right): Cc_NLP1.1 (CC_29_g12228), Cc_NLP1.2A (CC_29_g13117), Cc_NLP1.2B (CC_29_g13473), and Cc_NLP1.3 (CC_29_g14222). Rows correspond to time in hours post inoculation (hpi) (from bottom to top): 0, 6, 12, and 18. Scale bar corresponds to fold variation. Heatmap generated based on Fig. S3.

Although expression of the effector gene CC_29_g13117 (Cc_NLP1.2A) remained always very low during the time-points evaluated, there was some significant increase in expression towards the end of the experiment, at 18 hpi. Meanwhile, the effector gene CC_29_g13473 (Cc_NLP1.2B) reached levels of expression that were intermediate between gene CC_29_g12228 (Cc_NLP1.1) and gene CC_29_g13117 (Cc_NLP1.2A), with levels of expression increasing from 12 hpi onwards. The non-effector NLP gene CC_29_g14222 (Cc_NLP1.3) showed low (almost undetectable) levels of expression during the time-points evaluated.

### Overview on the effector NLPs of *C. cassiicola*

Herein, we proposed a scheme to illustrate the generic protein structure of an effector NLP in *C. cassiicola* (Fig. 5A). The sizes of the effector proteins varied from 237 (Cc_NLP1.1) to 243 (Cc_NLP1.2B) amino acids long. The protein consisted of a signal peptide (predicted by SignalP) and a single NPP1 domain (predicted by HMMER, PfamScan, and InterProScan). The sizes of the signal peptides varied from 18 (in Cc_NLP1.2A and Cc_NLP1.2B) to 21 (in Cc_NLP1.1) amino acids long. Sizes of the NPP1 domains (according to HMMER) varied from 185 (in Cc_NLP1.1) to 190 (in Cc_NLP1.2A and Cc_NLP1.2B) amino acids long. A segment varying from 27 (in Cc_NLP1.1) to 33 (in Cc_NLP1.2B) amino acids connected the signal peptide to the NPP1 domain. After the NPP1 domain, there were additional two (in Cc_NLP1.2A and Cc_NLP1.2B) to four (in Cc_NLP1.1) amino acids at the C-terminus of the protein. The two conserved cysteine residues that play crucial role in protein stabilization were located at the N-terminal side of the NPP1 domain. The positively selected sites that *codeml*/PAML analysis had identified were also located in the initial part of NPP1 domain of the effector NLPs of the Cc_NLP1.2B haplogroup.

**Fig. 5.**
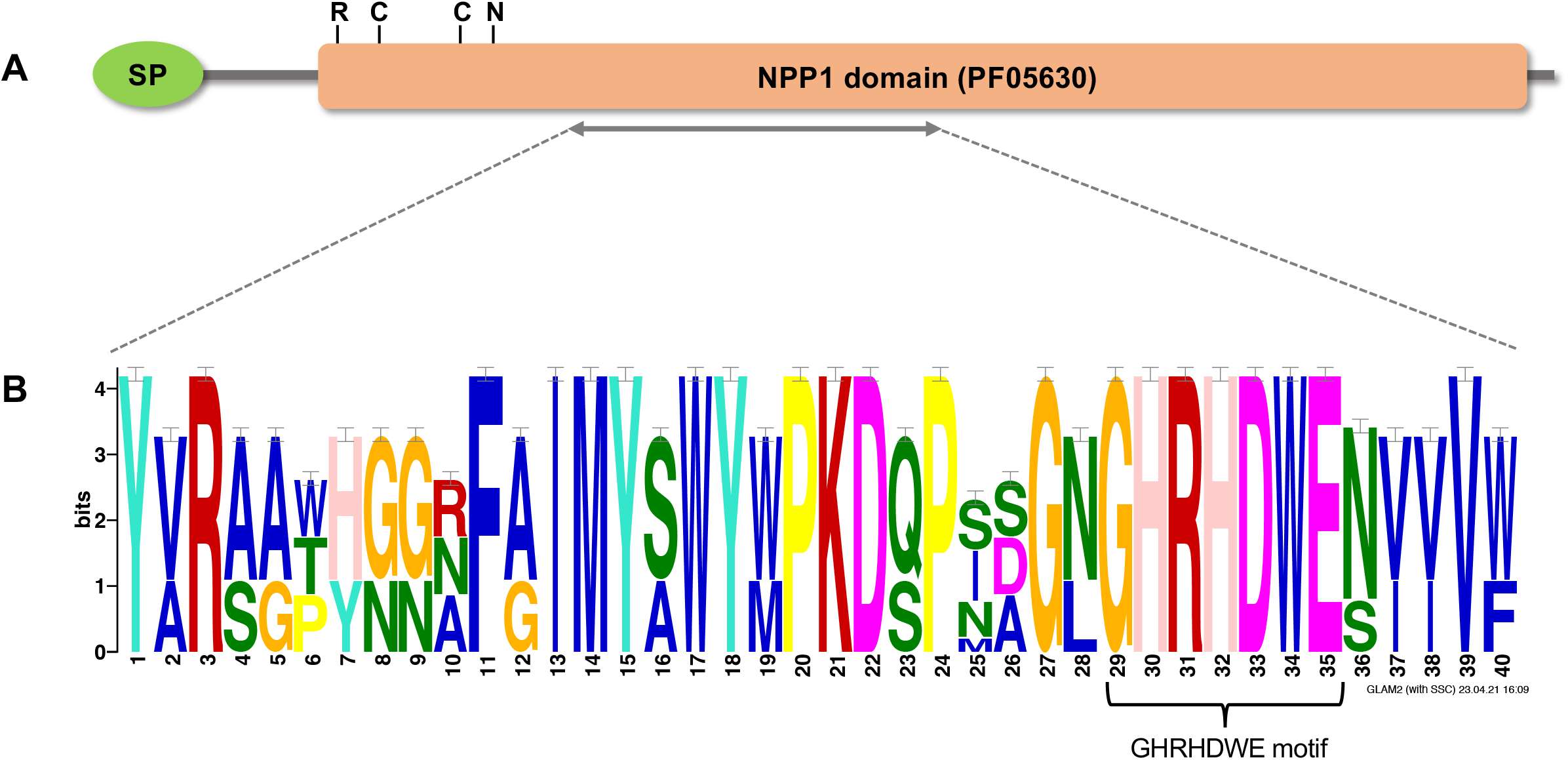
Overview of the Necrosis- and Ethylene-inducing peptide 1-like (NLP) effector proteins of *Corynespora cassiicola*. **A**. SignalP predicted the signal peptide (SP); HMMER predicted the NPP1 domain (PF05630). Horizontal grey segments depict the remaining amino acid residues. Over the NPP1 domain: C, conserved cysteine residues; R36 and N77, two positively selected residues of the Cc_NLP1.2B effector (see Fig. 4). **B**. Consensus pattern within the three NLP effectors (Cc_NLP1.1, Cc_NLP1.2A, and Cc_NLP1.2B). The variation pattern of amino acids surrounding the conserved GHRHDWE motif (positions 29-35) was generated with GLAM2SCAN and was visualized with GLAM2. Positions 1-3, 11-24, and 36-40 depict semiconserved sites. Position 22 contains a conserved residue of aspartic acid that is directly involved in cation binding.

A consensus pattern for the three effector NLPs (Cc_NLP1.1, Cc_NLP1.2A, and Cc_NLP1.2B) revealed the conserved GHRHDWE motif in the central region of the protein (positions 29-35, Fig. 5B). This conserved motif was surrounded by two semiconservative sites (positions 11-24 and 36-40). Together, this semiconserved region was part of two β-strands in the central portion of the protein (Fig. 4). One residue involved in the cation binding (aspartic acid, D) was conserved among the three effector NLPs (position 22, Fig. 5B) in *C. cassiicola* and was located in the negatively charged (cation binding) cavity of the protein secondary structure (Fig. 3).

The non-effector Cc_NLP1.3 is interesting. It diverged greatly in amino acid sequence from the counterpart NLPs (Cc_NLP1.1, Cc_NLP1.2A, and Cc_NLP1.2B) that were considered to be effectors. The highest level of sequence similarity was found between Cc_NLP1.3 and Cc_NLP1.2B (49%) and the lowest between Cc_NLP1.3 and Cc_NLP1.1 (30%). This higher level of similarity between Cc_NLP1.3 and Cc_NLP1.2B was congruent with the placement of Cc_NLP1.3 in the phylogenetic analysis (Fig. 1) and network analysis (Fig. 2). In Cc_NLP1.3, there was a tyrosine (Y) at the fourth residue of the heptapeptide motif (GHRYDWE) instead of the histidine (H) found in the motif of effector NLPs (GHRHDWE, Fig. 5B). Moreover, the NPP1 domain of Cc_NLP1.3 was shorter compared to those predicted for the effector NLPs: it consisted of 109 amino acids and started immediately from the GHRYDWE motif. Thus, semiconserved sites located before the heptapeptide motif in effector NLPs were not present in the non-effector Cc_NLP1.3 of *C. cassiicola*.

## DISCUSSION

### The NLPs of *C. cassiicola*

The likely role of NLPs as virulence factors during the early stages of infection and disease development (Fellbrich et al. 2002; Gijzen and Nürnberger, 2006; Levin et al. 2019; Pour et al. 2020; Duhan et al. 2021), the previous finding that the NLPs comprised a small gene family (Dal’Sasso et al 2022), and the worldwide emergence of target spot in economically important crops (Lopez et al. 2018; Sumabat et al. 2018; Banguela-Castillo et al 2020; Gao et al. 2020;) instigated us into exploring the diversity of NLPs in the plant pathogen *C. cassiicola*.

The size of the NLP superfamily has been shown to be consistently small across the Dothideomycetes, (Dal’Sasso et al. 2022). Most of the 79 studied species presented a single NLP gene; except the Botryosphaeriales, in which the NLP superfamily reached up to six members (Dal’Sasso et al. 2022). Among the Pleosporales, most of the 38 studied species exhibited up to two NLP genes. *Corynespora cassiicola* (isolate CCP) was one of the few species of Pleosporales that exhibited three NLP members, all of which were predicted to be effector genes within the NLP1 family (Dal’Sasso et al. 2022). Herein, we confirmed that the size of the NLP gene family across 44 isolates of *C. cassiicola* is consistently small, but differs from the family size we anticipated. The NLP family of a set of few isolates possesses an additional, fourth NLP copy, which was predicted to be a non-effector gene.

Biased gene retention (Dal’Sasso et al. 2022) was a likely force that maintained the three NLPs effector-encoding genes (Cc_NLP1.1, Cc_NLP1.2A, and Cc_NLP1.2B) in genomes of the 44 isolates of *C. cassiicola*. It is plausible that the maintenance of those genes contributed to facilitate early pathogen-host interaction. Likely, gene duplication from an NLP1.2 paralog originated Cc_NLP1.3, a forth NLP gene present in three isolates that were obtained from soybean. The Cc_NLP1.3 gene was predicted to be a non-effector gene; it accumulated many exclusive mutations and diverged greatly from other NLP encoding-genes, up to the point of losing several nucleotides that would encode amino acid residues shown to be required for protein functionality. This scenario suggests the Cc_NLP1.3 gene may be undergoing a process of pseudogenization as has been suggested previously for other NLP genes (Dal’Sasso et al. 2022) and supports the idea that effector genes undergo rapid turnover rates (the rate at which genes are duplicated and deleted) (Menardo et al. 2017). Moreover, the large number of mutational steps separating the four NLP haplogroups and the high levels of nucleotide diversity within each of those haplogroups are congruent with fungal effectors that undergo rapid evolution (Lo Presti et al. 2015; Menardo et al. 2017; Sánchez-Vallet et al. 2018).

### Selection upon the NLP effector–encoding genes

The three NLP effector-encoding genes of *C. cassiicola* were under different selective constraints. Although the intraspecific selective neutrality tests (Tajima’s D, Fu and Li’s D* and F*) indicated that the Cc_NLP1.1 gene was evolving under neutrality, the relative lower levels of nucleotide diversity and the nature/distribution of polymorphisms along the Cc_NLP1.1 haplotypes suggest that this locus was under stronger selective constraints, with deleterious polymorphisms being purged. The NLP genes homologous to the Cc_NLP1.1 exhibited widespread distribution among Dothideomycetes (Dal’Sasso et al. 2022). In species with multiple NLPs, the Cc_NLP1.1 homologues were the ones that showed the highest levels of cytotoxic to plants (Bashi et al. 2010; Fang et al. 2017; Levin et al. 2019; Pour et al. 2020; Dal’Sasso et al. 2022).

The large number of mutations in the Cc_NLP1.2A gene suggests that it is evolving under more relaxed selective constraints when compared to the others effector NLPs of *C. cassiicola*. Negative values of Fu and Li’s D* and F* together with the large number of low frequency haplotypes and missing intermediates suggest that the Cc_NLP1.2A gene has undergone a recent expansion. Accumulation of mutations as a source of genetic variability exposes effector genes to mutations with both beneficial and deleterious effects (Fouché et al. 2018). Rapid evolution leading to an excess of mutations could eventually lead to loss-of-function of the Cc_NLP1.2A in *C. cassiicola* and would help to explain the rare occurrence of Cc_NLP1.2A homologues across the Dothideomycetes (Dal’Sasso et al. 2022).

Selection analyses indicate that both balancing and positive selection are acting upon Cc_NLP1.2B. Significant positive values estimated by selective neutrality tests (Fu and Li’s D* and F*) indicates that balancing selection is acting to maintain the high levels of genetic variation among Cc_NLP1.2B haplotypes. Moreover, effector proteins under positive selection are often considered to operate in the frontline of host-pathogen interactions (Lo Presti et al. 2015). The finding that there were significant signals of positive selection acting upon the Cc_NLP1.2B gene agrees well with previous studies that identified positive selection in few (but not all) NLP genes of *Botrytis* spp. (Staats et al. 2007) and *Phytophthora sojae* (Dong et al. 2012). Furthermore, the amino acid residues that corresponded to N77 of the Cc_NLP1.2B protein are also under positive selection in a few effector NLPs from *P. sojae* (Dong et al. 2012). The biological implication of such positively selected sites remains elusive.

Even thought the effector NLPs of *C. cassiicola* diverged from each other in their amino acid sequences, they maintained a semiconserved central segment of amino acids among its effector NLP members (Fig. 5B). Previous reports have demonstrated that this semiconserved segment is the part of NLPs responsible for inducing typical MAMP-triggered defense responses in plants (Böhm et al. 2014; Oome et al. 2014). Therefore, purifying selection is likely acting upon NLP effector-encoding genes to maintain those key residues free of changes.

### Expression patterns of NLP genes during early soybean infection

Expression analyses confirmed that the NLPs are part of the effector armory that *C. cassiicola* uses during the early hours of infection. The effector NLPs exhibited different levels of gene expression. Although protein sequence analyses have indicated that all three effector NLPs of *C. cassiicola* were potentially functional, expression profiles revealed that significant expression at the early hours of soybean colonization took place for the Cc_NLP1.1 gene only. A gene homologous to Cc_NLP1.1 also was differentially expressed when the isolate CCP colonized rubber leaves (Lopez et al. 2018). In many plant pathogenic fungi, the genes that are homologues to Cc_NLP1.1 also exhibited high expression levels and strong cytotoxicity activities in the host cells (Bashi et al. 2010; Fang et al. 2017; Levin et al. 2019; Pour et al. 2020).

The low, but increasing levels of gene expression of the Cc_NLP1.2B gene may be due to a time-increasing gene expression cascade. It agrees with the hypothesis of redundancy of NLP copies as a strategy of functional diversification to overcome the host surveillance in a time-dependent manner (Seidl et al. 2015; Duhan et al. 2021; Dal’Sasso et al. 2022). The NLP encoding-genes are usually expressed in the early hours of host colonization but their expression could follow up to days, depending on the time needed by the pathogen to penetrate the host tissue (Levin et al. 2019; Pour et al. 2020; Duhan et al. 2021). Taken together, our gene expression analyses suggested that the expression of the NLP effector genes may occur in different levels during host colonization; likely, not all NLP effector genes are essential for pathogenicity of *C. cassiicola* to soybean.

## Acknowledgement

We thank Dr. Eveline Caixeta and the Laboratório de Biotecnologia do Cafeeiro for the use of the Real-Time PCR System. This work was supported by The Minas Gerais State Foundation of Research Aid – FAPEMIG (grant number APQ-00150-17) and by The National Council of Scientific and Technological Development – CNPq (fellowship number PQ 302336/2019-2) to LOO. TCSD received student fellowships from the CAPES Foundation (PROEX – 0487 No. 1684083) and CNPq (GM/GD 142400/2018-1).

## Supplementary material

**Fig. S1.**
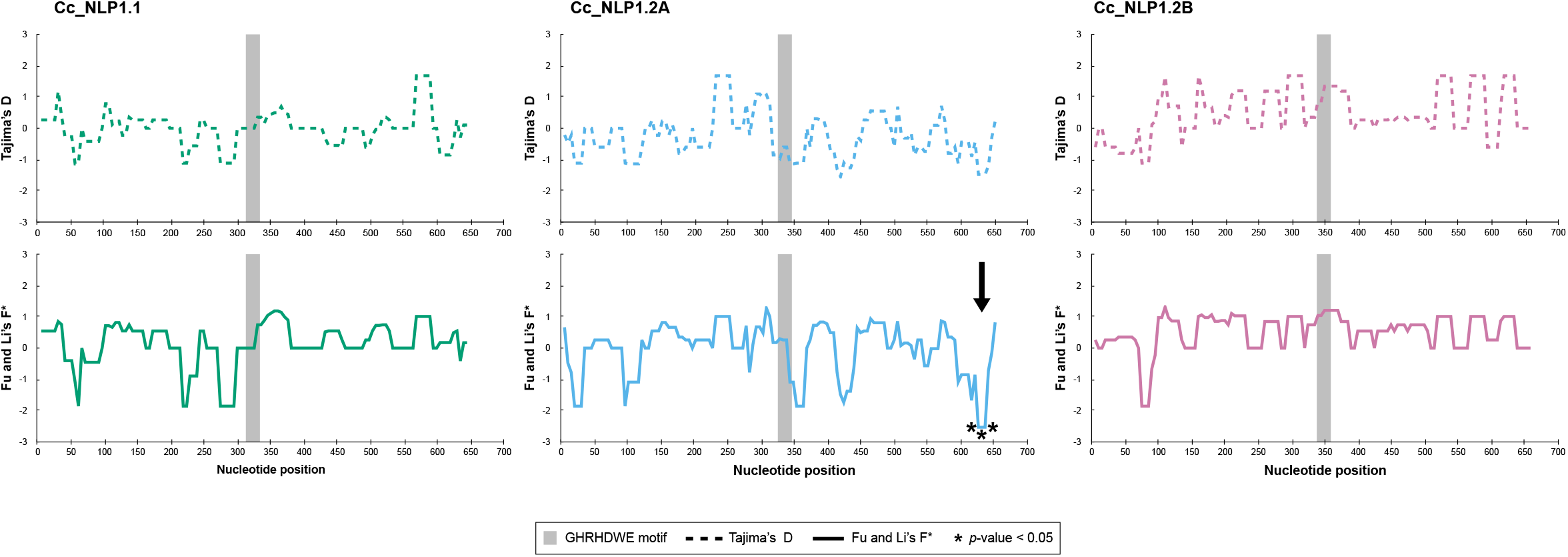
Sliding window neutrality tests of three predicted Necrosis- and Ethylene-inducing peptide 1-like protein (NLP)-encoding genes (CDS) of *Corynespora cassiicola*. The x axis shows the midpoints of contiguous windows (25 bases, with steps of 5 bases). The grey vertical bar indicates the relative position of the GHRHDWE motif. The arrow indicates sites with significant departures from zero (*p*-value < 0.05).

**Fig. S2.**
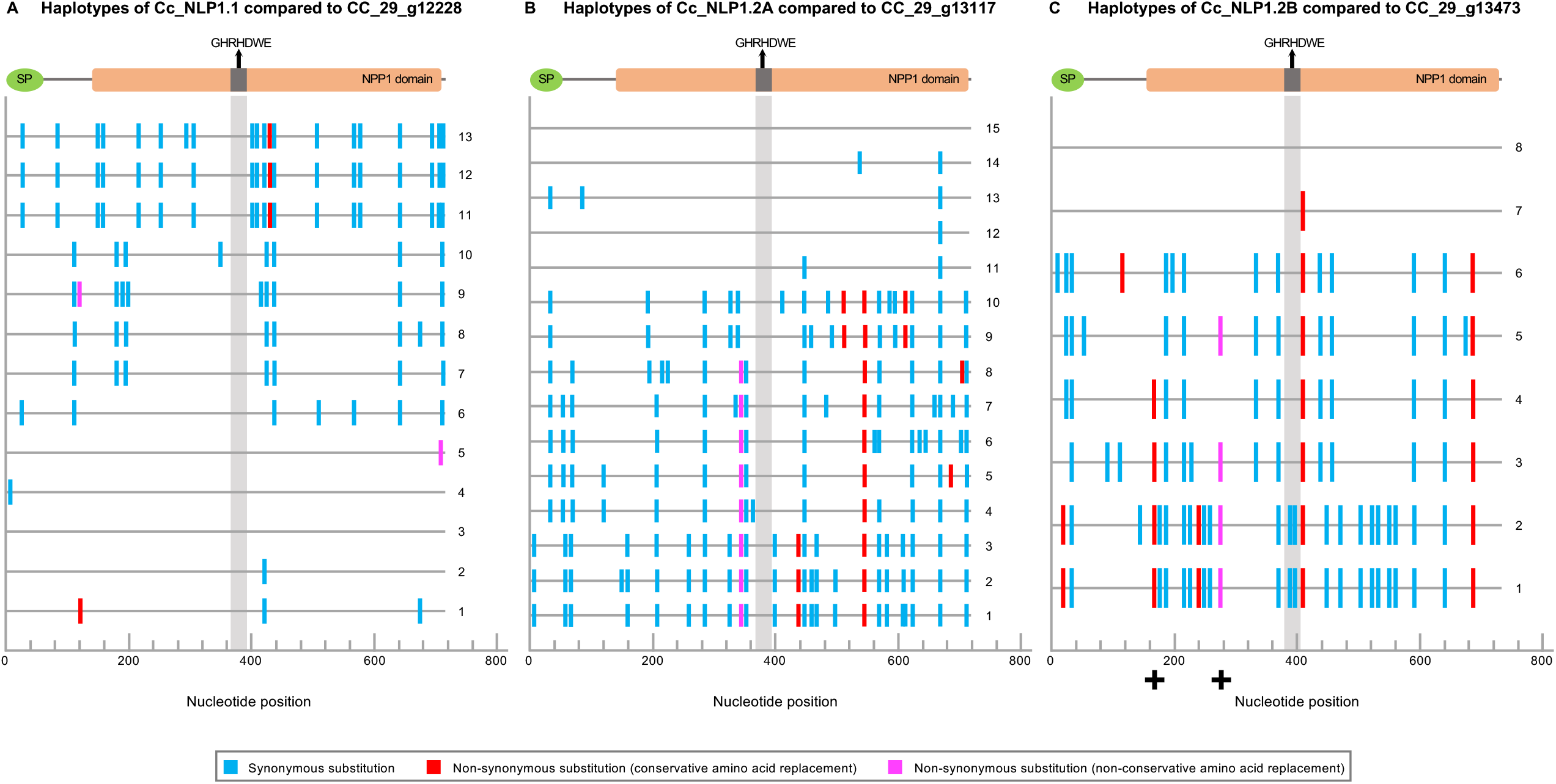
Placement of nucleotide substitutions along the haplotypes of each of the three NLP effector-encoding genes (Cc_NLP1.1, Cc_NLP1.2A, and Cc_NLP1.2B) (CDS) in *Corynespora cassiicola*. The NLP effector-encoding genes from isolate CC_29 were used as reference for haplotype sequence alignment, according to the haplogroups in Fig. 3. Upper protein representations indicate portions of the CDS that encode for the signal peptide (SP), the NPP1 domain, and the conserved GHRHDWE motif in the effector NLPs of isolate CC_29. **A**. Cc_NLP1.1 compared to CC_29_g12228 (haplotype 3). **B**. Cc_NLP1.2A compared to CC_29_g13117 (haplotype 15). **C**. Cc_NLP1.2B compared to CC_29_g13473 (haplotype 8). Plus (+) signs indicate substitutions that resulted in amino acids replacements predicted to be positively selected, according to *codeml*/PAML analysis. Types of nucleotide substitutions as indicated.

**Fig. S3.**
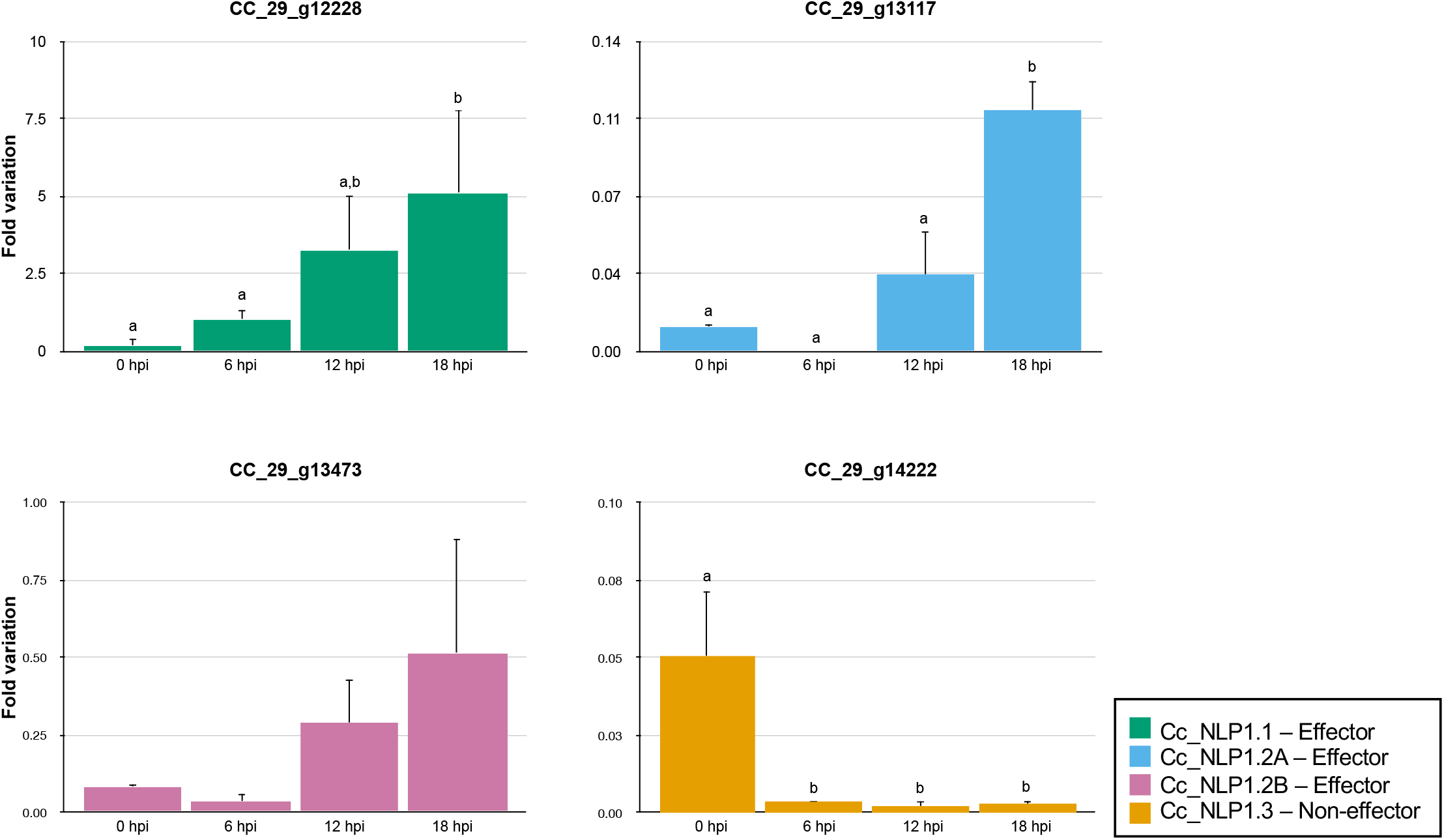
Expression patterns of the Necrosis- and Ethylene-inducing peptide 1-like (NLP) genes of *Corynespora cassiicola* (isolate CC_29) after inoculation in soybean plants. Gene expression was inferred by RT-qPCR analyses. The time (x axis) is given in hours post inoculation (hpi). The fold variation of gene expression of each NLP gene (y axis) was calculated using the 2^-ΔCt^ method, the values were normalized to an endogenous control gene (β-tubulin). Letters show significant differences (*p*-value < 0.05) after a one-way ANOVA followed by Tukey’s test for each NLP gene individually. Error bars represent the standard deviation of three biological replicates.

**TableS1**. Detailed information about the isolates of *Corynespora cassiicola* used in this study.

**Table S2**. Primers designed for *Corynespora cassiicola* isolate CC_29 for NLP gene expression analyses using RT-qPCR.

**Table S3**. Summary statistics on *de novo* genome assemblies of *Corynespora cassiicola* isolates India_Hevea and TScotton1.

