## Supplementary material for "The Necrosis- and Ethylene-inducing peptide 1-like protein (NLP) gene family of the plant pathogen *Corynespora cassiicola*": Table S1

**TableS1.** Detailed information on all isolates of *Corynespora cassiicola* used in this study.

| Isolate | BioSample | BioProject | Host Species | Geographic Origin | Genome Size (Mb) | # Genes | Reference | This work | | | | |
|---|---|---|---|---|---|---|---|---|---|---|---|---|
| | | | | | | | | # NLP Genes | # NLP Effector Genes | NLP Gene ID | Haplogroup | Haplotype |
| 777AA | SAMN08330686 | PRJNA428435 | *Glycine max* | Brazil | 41.70 | 14,873 | Lopez et al. (2018) | 4 | 3 | 777AA_g10179 | Cc_NLP1.1 | 2 |
| | | | | | | | | | | 777AA_g880 | Cc_NLP1.2A | 11 |
| | | | | | | | | | | 777AA_g10197 | Cc_NLP1.2B | 8 |
| | | | | | | | | | | 777AA_g14294 | Cc_NLP1.3 | 1 |
| ATI11 | SAMN08330687 | PRJNA428435 | *Cucumis sativus* | Brazil | 41.13 | 14,497 | Lopez et al. (2018) | 3 | 3 | ATI11_g10166 | Cc_NLP1.1 | 3 |
| | | | | | | | | | | ATI11_g1730 | Cc_NLP1.2A | 15 |
| | | | | | | | | | | ATI11_g11360 | Cc_NLP1.2B | 8 |
| C7_L008 | SAMEA103891068 | PRJEB19843 | - | - | 42.50 | 14,569 | - | 3 | 3 | C7_L008_g6805 | Cc_NLP1.1 | 4 |
| | | | | | | | | | | C7_L008_g10130 | Cc_NLP1.2A | 11 |
| | | | | | | | | | | C7_L008_g6820 | Cc_NLP1.2B | 7 |
| CAL-4 | SAMN12086008 | PRJNA549429 | *Gossypium hirsutum* | United States | 44.90 | 19,240 | - | 3 | 3 | CAL-4_g12966 | Cc_NLP1.1 | 2 |
| | | | | | | | | | | CAL-4_g6936 | Cc_NLP1.2A | 11 |
| | | | | | | | | | | CAL-4_g13172 | Cc_NLP1.2B | 8 |
| CBS 129. | SAMN08330688 | PRJNA428435 | *Cucumis sativus* | Brazil | 40.79 | 14,403 | Lopez et al. 2018 | 3 | 3 | CBS12925_g6790 | Cc_NLP1.1 | 2 |
| | | | | | | | | | | CBS12925_g3518 | Cc_NLP1.2A | 15 |
| | | | | | | | | | | CBS12925_g6808 | Cc_NLP1.2B | 8 |
| CC_29 | SAMN18718492 | PRJNA721456 | *Glycine max* | Brazil | 44.79 | 18,487 | Dal'Sasso et al. 2021 | 4 | 3 | CC_29_g12228 | Cc_NLP1.1 | 3 |
| | | | | | | | | | | CC_29_g13117 | Cc_NLP1.2A | 15 |
| | | | | | | | | | | CC_29_g13473 | Cc_NLP1.2B | 8 |
| | | | | | | | | | | CC_29_g14222 | Cc_NLP1.3 | 3 |
| CCAM1 | SAMN08330689 | PRJNA428435 | *Hevea brasiliensis* | Cameroon | 42.33 | 14,634 | Lopez et al. 2018 | 3 | 3 | CCAM1_g9108 | Cc_NLP1.1 | 7 |
| | | | | | | | | | | CCAM1_g5347 | Cc_NLP1.2A | 4 |
| | | | | | | | | | | CCAM1_g13882 | Cc_NLP1.2B | 4 |
| CCAM2 | SAMN08330690 | PRJNA428435 | *Hevea brasiliensis* | Cameroon | 41.89 | 14,785 | Lopez et al. 2018 | 3 | 3 | CCAM2_g9764 | Cc_NLP1.1 | 11 |
| | | | | | | | | | | CCAM2_g3518 | Cc_NLP1.2A | 6 |
| | | | | | | | | | | CCAM2_g10878 | Cc_NLP1.2B | 5 |
| CCAM3 | SAMN08330691 | PRJNA428435 | *Hevea brasiliensis* | Cameroon | 42.72 | 15,022 | Lopez et al. 2018 | 3 | 3 | CCAM3_g102 | Cc_NLP1.1 | 11 |
| | | | | | | | | | | CCAM3_g10005 | Cc_NLP1.2A | 6 |
| | | | | | | | | | | CCAM3_g12335 | Cc_NLP1.2B | 5 |
| CCAM4 | SAMN08330692 | PRJNA428435 | *Hevea brasiliensis* | Cameroon | 42.32 | 14,545 | Lopez et al. 2018 | 3 | 3 | CCAM4_g6615 | Cc_NLP1.1 | 7 |
| | | | | | | | | | | CCAM4_g3778 | Cc_NLP1.2A | 4 |
| | | | | | | | | | | CCAM4_g9511 | Cc_NLP1.2B | 4 |
| CCI13 | SAMN08330693 | PRJNA428435 | *Hevea brasiliensis* | Côte d'Ivoire | 41.94 | 14,847 | Lopez et al. 2018 | 3 | 3 | CCI13_g7758 | Cc_NLP1.1 | 3 |
| | | | | | | | | | | CCI13_g2701 | Cc_NLP1.2A | 11 |
| | | | | | | | | | | CCI13_g12991 | Cc_NLP1.2B | 7 |
| CCI6 | SAMN08330694 | PRJNA428435 | *Hevea brasiliensis* | Côte d'Ivoire | 41.99 | 14,905 | Lopez et al. 2018 | 3 | 3 | CCI6_g6508 | Cc_NLP1.1 | 3 |
| | | | | | | | | | | CCI6_g12011 | Cc_NLP1.2A | 11 |
| | | | | | | | | | | CCI6_g10427 | Cc_NLP1.2B | 7 |
| CCP | SAMN05660679 | PRJNA234811 | *Hevea brasiliensis* | Philippines | 44.85 | 17,166 | Lopez et al. 2018 | 3 | 3 | CCP_601720 | Cc_NLP1.1 | 11 |
| | | | | | | | | | | CCP_537044 | Cc_NLP1.2A | 6 |
| | | | | | | | | | | CCP_497672 | Cc_NLP1.2B | 5 |
| CGAB1 | SAMN08330695 | PRJNA428435 | *Hevea brasiliensis* | Gabon | 41.20 | 14,635 | Lopez et al. 2018 | 3 | 3 | CGAB1_g7423 | Cc_NLP1.1 | 3 |
| | | | | | | | | | | CGAB1_g5586 | Cc_NLP1.2A | 14 |
| | | | | | | | | | | CGAB1_g8330 | Cc_NLP1.2B | 7 |
| CGAB2 | SAMN08330696 | PRJNA428435 | *Hevea brasiliensis* | Gabon | 41.22 | 14,638 | Lopez et al. 2018 | 3 | 3 | CGAB2_g869 | Cc_NLP1.1 | 3 |
| | | | | | | | | | | CGAB2_g1004 | Cc_NLP1.2A | 14 |
| | | | | | | | | | | CGAB2_g3883 | Cc_NLP1.2B | 7 |
| CIND3 | SAMN08330697 | PRJNA428435 | *Hevea brasiliensis* | India | 41.67 | 14,733 | Lopez et al. 2018 | 3 | 3 | CIND3_g13589 | Cc_NLP1.1 | 3 |
| | | | | | | | | | | CIND3_g13380 | Cc_NLP1.2A | 11 |
| | | | | | | | | | | CIND3_g14081 | Cc_NLP1.2B | 7 |
| CLN16 | SAMN08330698 | PRJNA428435 | *Hevea brasiliensis* | Malaysia | 41.71 | 14,803 | Lopez et al. 2018 | 3 | 3 | CLN16_g9268 | Cc_NLP1.1 | 3 |
| | | | | | | | | | | CLN16_g9820 | Cc_NLP1.2A | 11 |

| | | | | | | | | | | | | |
|---|---|---|---|---|---|---|---|---|---|---|---|---|
| | | | | | | | | | | CLN16_g10739 | Cc_NLP1.2B | 7 |
| CSB16 | SAMN08330699 | PRJNA428435 | *Hevea brasiliensis* | Malaysia | 40.65 | 14,489 | Lopez et al. 2018 | 3 | 3 | CSB16_g4822 | Cc_NLP1.1 | 12 |
| | | | | | | | | | | CSB16_g11473 | Cc_NLP1.2A | 9 |
| | | | | | | | | | | CSB16_g4841 | Cc_NLP1.2B | 1 |
| CSRI1 | SAMN08330700 | PRJNA428435 | *Hevea brasiliensis* | Sri Lanka | 40.61 | 14,327 | Lopez et al. 2018 | 3 | 3 | CSRI1_g7251 | Cc_NLP1.1 | 12 |
| | | | | | | | | | | CSRI1_g13432 | Cc_NLP1.2A | 9 |
| | | | | | | | | | | CSRI1_g7232 | Cc_NLP1.2B | 1 |
| CSRI2 | SAMN08330701 | PRJNA428435 | *Hevea brasiliensis* | Sri Lanka | 40.77 | 14,391 | Lopez et al. 2018 | 3 | 3 | CSRI2_g9601 | Cc_NLP1.1 | 12 |
| | | | | | | | | | | CSRI2_g8039 | Cc_NLP1.2A | 9 |
| | | | | | | | | | | CSRI2_g9582 | Cc_NLP1.2B | 1 |
| CSRI5 | SAMN08330702 | PRJNA428435 | *Hevea brasiliensis* | Sri Lanka | 41.67 | 14,736 | Lopez et al. 2018 | 3 | 3 | CSRI5_g4338 | Cc_NLP1.1 | 3 |
| | | | | | | | | | | CSRI5_g10755 | Cc_NLP1.2A | 11 |
| | | | | | | | | | | CSRI5_g9398 | Cc_NLP1.2B | 7 |
| CTHA1 | SAMN08330703 | PRJNA428435 | *Hevea brasiliensis* | Thailand | 41.28 | 14,432 | Lopez et al. 2018 | 3 | 3 | CTHA1_g5888 | Cc_NLP1.1 | 9 |
| | | | | | | | | | | CTHA1_g12212 | Cc_NLP1.2A | 8 |
| | | | | | | | | | | CTHA1_g1711 | Cc_NLP1.2B | 6 |
| CTHA2 | SAMN08330704 | PRJNA428435 | *Hevea brasiliensis* | Thailand | 40.82 | 14,424 | Lopez et al. 2018 | 3 | 3 | CTHA2_g8146 | Cc_NLP1.1 | 12 |
| | | | | | | | | | | CTHA2_g8485 | Cc_NLP1.2A | 9 |
| | | | | | | | | | | CTHA2_g8165 | Cc_NLP1.2B | 1 |
| CTHA3 | SAMN08330705 | PRJNA428435 | *Hevea brasiliensis* | Thailand | 40.37 | 14,616 | Lopez et al. 2018 | 3 | 3 | CTHA3_g11953 | Cc_NLP1.1 | 9 |
| | | | | | | | | | | CTHA3_g5642 | Cc_NLP1.2A | 8 |
| | | | | | | | | | | CTHA3_g4093 | Cc_NLP1.2B | 6 |
| CTHA4 | SAMN08330706 | PRJNA428435 | *Hevea brasiliensis* | Thailand | 40.71 | 14,410 | Lopez et al. 2018 | 3 | 3 | CTHA4_g13560 | Cc_NLP1.1 | 9 |
| | | | | | | | | | | CTHA4_g2821 | Cc_NLP1.2A | 8 |
| | | | | | | | | | | CTHA4_g6803 | Cc_NLP1.2B | 6 |
| CTHA5 | SAMN08330707 | PRJNA428435 | *Hevea brasiliensis* | Thailand | 40.86 | 14,461 | Lopez et al. 2018 | 3 | 3 | CTHA5_g13106 | Cc_NLP1.1 | 5 |
| | | | | | | | | | | CTHA5_g688 | Cc_NLP1.2A | 11 |
| | | | | | | | | | | CTHA5_g13694 | Cc_NLP1.2B | 7 |
| CTHA6 | SAMN08330708 | PRJNA428435 | *Hevea brasiliensis* | Thailand | 40.94 | 14,444 | Lopez et al. 2018 | 3 | 3 | CTHA6_g4937 | Cc_NLP1.1 | 3 |
| | | | | | | | | | | CTHA6_g646 | Cc_NLP1.2A | 11 |
| | | | | | | | | | | CTHA6_g11768 | Cc_NLP1.2B | 7 |
| CVa5 | SAMN12086010 | PRJNA549429 | *Gossypium hirsutum* | United States | 44.52 | 15,348 | Lopez et al. 2018 | 3 | 3 | CVa5_g12316 | Cc_NLP1.1 | 2 |
| | | | | | | | | | | CVa5_g12873 | Cc_NLP1.2A | 11 |
| | | | | | | | | | | CVa5_g13757 | Cc_NLP1.2B | 8 |
| E139 | SAMN08330709 | PRJNA428435 | *Hevea brasiliensis* | Brazil | 42.82 | 14,648 | Lopez et al. 2018 | 3 | 3 | E139_g2766 | Cc_NLP1.1 | 8 |
| | | | | | | | | | | E139_g2931 | Cc_NLP1.2A | 7 |
| | | | | | | | | | | E139_g11927 | Cc_NLP1.2B | 4 |
| E55 | SAMN08330710 | PRJNA428435 | *Hevea brasiliensis* | Brazil | 40.53 | 14,352 | Lopez et al. 2018 | 3 | 3 | E55_g11737 | Cc_NLP1.1 | 6 |
| | | | | | | | | | | E55_g6497 | Cc_NLP1.2A | 2 |
| | | | | | | | | | | E55_g10462 | Cc_NLP1.2B | 3 |
| E79 | SAMN08330711 | PRJNA428435 | *Hevea brasiliensis* | Brazil | 42.47 | 14,373 | Lopez et al. 2018 | 3 | 3 | E79_g12376 | Cc_NLP1.1 | 10 |
| | | | | | | | | | | E79_g1196 | Cc_NLP1.2A | 5 |
| | | | | | | | | | | E79_g5219 | Cc_NLP1.2B | 4 |
| EDIG | SAMN08330712 | PRJNA428435 | *Cucumis sativus* | Brazil | 41.24 | 14,743 | Lopez et al. 2018 | 3 | 3 | EDIG_g11647 | Cc_NLP1.1 | 3 |
| | | | | | | | | | | EDIG_g11366 | Cc_NLP1.2A | 15 |
| | | | | | | | | | | EDIG_g6937 | Cc_NLP1.2B | 8 |
| GSO2 | SAMN08330713 | PRJNA428435 | *Vernonia cinerea* | Brazil | 41.80 | 15,072 | Lopez et al. 2018 | 3 | 3 | GSO2_g7213 | Cc_NLP1.1 | 3 |
| | | | | | | | | | | GSO2_g1195 | Cc_NLP1.2A | 12 |
| | | | | | | | | | | GSO2_g529 | Cc_NLP1.2B | 7 |
| IA | SAMN08330714 | PRJNA428435 | *Cucumis sativus* | Brazil | 41.58 | 14,793 | Lopez et al. 2018 | 3 | 3 | IA_g12863 | Cc_NLP1.1 | 2 |
| | | | | | | | | | | IA_g8583 | Cc_NLP1.2A | 15 |
| | | | | | | | | | | IA_g14565 | Cc_NLP1.2B | 8 |
| India_Hev | SAMN10434155 | PRJNA505742 | *Hevea brasiliensis* | India | 42.89 | 14,404 | - | 3 | 3 | India_Hevea_g13512 | Cc_NLP1.1 | 1 |
| | | | | | | | | | | India_Hevea_g4639 | Cc_NLP1.2A | 13 |
| | | | | | | | | | | India_Hevea_g13452 | Cc_NLP1.2B | 8 |
| JQ | SAMN08330715 | PRJNA428435 | *Cucumis sativus* | Brazil | 41.55 | 14,815 | Lopez et al. 2018 | 3 | 3 | JQ_g1090 | Cc_NLP1.1 | 2 |

| | | | | | | | | | | | |
|---|---|---|---|---|---|---|---|---|---|---|---|
| | | | | | | | | | JQ_g6266 | Cc_NLP1.2A | 15 |
| | | | | | | | | | JQ_g14090 | Cc_NLP1.2B | 8 |
| LPO7 | SAMN08330716 | PRJNA428435 | *Piper hispidinervum* | Brazil | 42.23 | 14,535 | Lopez et al. 2018 | 3 | 3 | LPO7_g9249 | Cc_NLP1.1 | 6 |
| | | | | | | | | | LPO7_g5694 | Cc_NLP1.2A | 1 |
| | | | | | | | | | LPO7_g9266 | Cc_NLP1.2B | 3 |
| PB | SAMN08330717 | PRJNA428435 | *Cucumis sativus* | Brazil | 41.56 | 14,801 | Lopez et al. 2018 | 3 | 3 | PB_g7351 | Cc_NLP1.1 | 2 |
| | | | | | | | | | PB_g3886 | Cc_NLP1.2A | 15 |
| | | | | | | | | | PB_g2314 | Cc_NLP1.2B | 8 |
| RUD | SAMN08330718 | PRJNA428435 | *Glycine max* | Brazil | 41.63 | 14,858 | Lopez et al. 2018 | 4 | 3 | RUD_g9099 | Cc_NLP1.1 | 3 |
| | | | | | | | | | RUD_g11509 | Cc_NLP1.2A | 15 |
| | | | | | | | | | RUD_g10304 | Cc_NLP1.2B | 8 |
| | | | | | | | | | RUD_g11851 | Cc_NLP1.3 | 2 |
| SS1 | SAMN08330719 | PRJNA428435 | *Hevea brasiliensis* | Malaysia | 40.73 | 14,378 | Lopez et al. 2018 | 3 | 3 | SS1_g6084 | Cc_NLP1.1 | 12 |
| | | | | | | | | | SS1_g6638 | Cc_NLP1.2A | 9 |
| | | | | | | | | | SS1_g6065 | Cc_NLP1.2B | 1 |
| TCI3 | SAMN12086019 | PRJNA549429 | *Solanum lycopersicum* | United States | 45.34 | 14,987 | - | 3 | 3 | TCI3_g7271 | Cc_NLP1.1 | 6 |
| | | | | | | | | | TCI3_g7342 | Cc_NLP1.2A | 3 |
| | | | | | | | | | TCI3_g12718 | Cc_NLP1.2B | 3 |
| TSB1 | SAMN08330720 | PRJNA428435 | *Hevea brasiliensis* | Malaysia | 40.89 | 14,435 | Lopez et al. 2018 | 3 | 3 | TSB1_g6254 | Cc_NLP1.1 | 12 |
| | | | | | | | | | TSB1_g3780 | Cc_NLP1.2A | 9 |
| | | | | | | | | | TSB1_g6273 | Cc_NLP1.2B | 1 |
| TScotton1 | SAMN06706475 | PRJNA382361 | *Gossypium hirsutum* | United States | 42.10 | 14,988 | Shrestha et al. 2018 | 3 | 3 | TScotton1_g11067 | Cc_NLP1.1 | 2 |
| | | | | | | | | | TScotton1_g11728 | Cc_NLP1.2A | 11 |
| | | | | | | | | | TScotton1_g11410 | Cc_NLP1.2B | 8 |
| UM591 | SAMN02981578 | PRJNA236064 | *Homo sapiens* | Malaysia | 41.35 | 14,695 | Looi et al. 2017 | 3 | 3 | UM591_g2079 | Cc_NLP1.1 | 13 |
| | | | | | | | | | UM591_g4896 | Cc_NLP1.2A | 10 |
| | | | | | | | | | UM591_g2097 | Cc_NLP1.2B | 2 |
