## Supplementary material for "The Necrosis- and Ethylene-inducing peptide 1-like protein (NLP) gene family of the plant pathogen *Corynespora cassiicola*": Table S2

**Table S2.** Primers designed for *Corynespora cassiicola* isolate CC_29 for NLP gene expression analyses using RT-qPCR.

| Gene Name | Gene ID | Forward sequence (5' – 3') | Reverse sequence (5' – 3') |
|---|---|---|---|
| Cc_NLP1.1 | CC_29_g12228 | CCTGTCGAATCCTCCCTTGA | AACGACCGATAGCACCACTC |
| Cc_NLP1.2A | CC_29_g13117 | GAAGTTTGCGCCCTACATGC | GAGTCCACCGCTCGTGTTTC |
| Cc_NLP1.2B | CC_29_g13473 | GGGTTCCCTCAGACTGTTCC | ATGGCACGCAACCATTGAAG |
| Cc_NLP1.3 | CC_29_g14222 | CCATTGGCAGGGCTATGAAGA | CCAGCATCATCCTTAGCGGT |
| $\beta$-tubulin | CC_29_g4872 | CAGACCGGTCAATGCGGTAA | GCCATTGTAGACGCCGGATC |
