## Supplementary material for "The Necrosis- and Ethylene-inducing peptide 1-like protein (NLP) gene family of the plant pathogen *Corynespora cassiicola*": Table S3

**Table S3.** Summary statistics on *de novo* genome assemblies of *Corynespora cassiicola* isolates India_Hevea and TScotton1.

| **Genomic statistics** | **India_Hevea** | **TScotton1** |
|---|---:|---:|
| **Assembly** | | |
| Total assembly size (Mbp) | 42.89 | 42.11 |
| # of scaffolds (>= 500 bp) | 1,059 | 2,188 |
| # of scaffolds (>= 10,000 bp) | 443 | 806 |
| # of scaffolds (>= 50,000 bp) | 246 | 282 |
| Largest scaffold (bp) | 827,165 | 251,612 |
| GC (%) | 50.85 | 52.56 |
| $N_{50}$ (bp) | 149,160 | 62,602 |
| Total reads | 21,333,739 | 74,123,165 |
| Mapped reads (%) | 99.80 | 99.54 |
| Average coverage depth (X) | 74 | 263 |
| **Gene completeness** | | |
| Complete BUSCOs (%) | 99.6 | 98.9 |
| Complete and single-copy BUSCOs (%) | 99.5 | 98.8 |
